# Deciphering molecular heterogeneity and dynamics of human hippocampal neural stem cells at different ages and injury states

**DOI:** 10.1101/2023.05.15.540723

**Authors:** Junjun Yao, Shaoxing Dai, Ran Zhu, Ju Tan, Qiancheng Zhao, Yu Yin, Jiansen Sun, Xuewei Du, Longjiao Ge, Jianhua Xu, Chunli Hou, Nan Li, Jun Li, Weizhi Ji, Chuhong Zhu, Runrui Zhang, Tianqing Li

## Abstract

While accumulated publications support the existence of neurogenesis in the adult human hippocampus, the homeostasis and developmental potentials of neural stem cells (NSCs) under different contexts remain unclear. Based on our generated single-nucleus atlas of the human hippocampus across neonatal, adult, aging and injury, we dissected the molecular heterogeneity and transcriptional dynamics of human hippocampal NSCs under different contexts. We further identified new specific neurogenic lineage markers that overcome the lack of specificity found in some well-known markers. Based on developmental trajectory and molecular signatures, we found that a subset of NSCs exhibit quiescent properties after birth, and most NSCs become deep quiescence during aging. Furthermore, certain deep quiescent NSCs are re-activated following stroke injury. Together, our findings provide valuable insights into the development, aging, and re-activation of the human hippocampal NSCs, and help to explain why adult hippocampal neurogenesis is infrequently observed in humans.

## Introduction

Continuous learning and memory formation throughout life is driven by developmental and adult neurogenesis. The dentate gyrus (DG), a part of the hippocampus and one of the main neurogenic niches, sustains neurogenesis through the activity of resident neural stem cells (NSCs) (Berg et al. 2019). Although adult neurogenesis in rodents (Hochgerner et al. 2018; Dulken et al. 2017) is well studied, and age-related neurogenesis decline is conserved across species, whether hippocampal neurogenesis persists in the adult human brain has been debated over the years. Finding a conclusive answer to this question is not trivial, as available human brain tissue is rare, and analysis is fraught with technical challenges. Based on marker immunostaining, a few studies found no evidence of neurogenesis in human after adolescence (Cipriani et al. 2018; Sorrells et al. 2018; Franjic et al. 2022), while others detected that human neurogenesis persists in adulthood but declines during aging (Boldrini et al. 2018; Moreno-Jimenez et al. 2019; Tobin et al. 2019; Terreros-Roncal et al. 2021). It is expected that single-cell RNA-sequencing will help resolve the ongoing debate, as this technology is capable of bypassing the biases associated with traditional methods of immunostaining and quantification. Single cell analysis approaches can also help identify novel cell markers and resolve the dynamics of transcriptional signatures during neurogenesis under different conditions. Leveraging these advantages, several groups performed single-nucleus RNA-seq (snRNA-seq) (Habib et al. 2016) analysis to investigate adult hippocampal neurogenesis in the human brain (Franjic et al. 2022; Zhou et al. 2022; Wang et al. 2022b). Although one study failed to detect evidence of adult neurogenic trajectories in human hippocampal tissues (Franjic et al. 2022), the other two reported the presence of molecular programs consistent with the capacity for the adult human dentate gyrus to generate new granule cells (Zhou et al. 2022; Wang et al. 2022b).

Accumulated publications support the existence of neurogenesis in the adult human hippocampus, but the homeostasis and developmental potentials of NSCs under different contexts remain unclear. Particularly, while actively proliferating in early development, mouse NSCs gradually acquire quiescent properties and transform into quiescent NSCs (qNSCs) with age. Although neurogenesis declines in the mouse aging hippocampus as a consequence of NSC loss and dormancy, qNSCs can be reactivated into active NSCs (aNSCs) that give rise to granule cells (GCs) which integrate into existing neural circuits (Encinas et al. 2011; Obernier and Alvarez-Buylla 2019). Specifically, ischemic insult in the adult mouse brain has been reported to evoke qNSC to transition into an active state. However, whether these similar mechanisms occur in the human hippocampus is unknown (E. Llorens-Bobadilla et al. 2015a).

To gain insight into why adult hippocampal neurogenesis is challenging to observe in humans, we believe that examining NSCs under varying conditions may be helpful, as they are the source of neurogenesis. Thus, we conducted snRNA-seq analysis on human hippocampal tissue and investigated the heterogeneity and molecular dynamics of hippocampal NSCs across neonatal, adult, aging and stroke-induced injury conditions. Based on comparative analysis of cell types, and developmental trajectories and molecular features of NSCs under different contexts, we found that NSCs, including qNSCs, primed NSCs (pNSCs) and aNSCs, exhibit different molecular features and dynamics across neonatal, adult, aging, and stroke-induced injury conditions. We observed a subset of NSCs that display quiescent properties after birth, and most NSCs become deep quiescence during aging. Notably, some deep qNSCs can be reactivated to give rise to pNSCs and aNSCs in the stroke-injured adult human hippocampus. In addition, we also found that immature granule cell markers widely used in mice studies, including DCX and PROX1, are non-specifically expressed in human hippocampal GABAergic interneurons. We further identified neuroblast-specific genes *CALM3, NEUROD2, NRN1* and *NRGN* with low/absent expression in human GABAergic interneurons. Together, our findings provide an important resource to understand the development, aging and activation of human postnatal hippocampal NSCs.

## Results

### Single-nucleus atlas of the human hippocampus across ages and injury

To generate a comprehensive cell atlas of neurogenic lineages in the human hippocampus, we collected 10 donated post-mortem hippocampal tissues. We then dissociated the anterior-mid hippocampus (which has an obvious dentate gyrus structure) and performed 10x genomics single-nucleus RNA-sequencing. We also performed immunostaining for the counterpart side of each hippocampus sample (**Figure 1A**). The 10 individual samples, divided into four groups according to age and brain health, included neonatal (Day 4 after birth, D4, n=1), adult (31-, 32-year-old, n=2), aging (from 50 to 68-year-old, n=6) and stroke-induced injury (48-year-old, n=1) groups (**Figure 1A, Figure 1-source data 1**). In total, we sequenced 99,635 single nuclei of which 92,966 nuclei were successfully retained after quality control and filtration. After the removal of cell debris, cell aggregates, and cells with more than 20% of mitochondrial genes transcripts, we analyzed a median of 3001 genes per nucleus. (**Figure1-figure supplement 1A and B**). To generate an overview of hippocampal cell types, we pooled single cells from all samples and categorized human hippocampal cells based on classical markers and differentially expressed genes (DEGs) into 16 main populations by Uniform Manifold Approximation and Projection (UMAP) (**Figure 1B to D, Figure1-figure supplement 1C to E**). These included astrocyte1 (AS1), astrocytes2/quiescent neural stem cell (AS2/qNSC), pNSC, aNSC, neuroblast (NB), granule cell (GC), GABAergic interneuron (GABA-IN), pyramidal neuron (PN), oligodendrocyte progenitor (OPC), oligodendrocyte (OLG), microglia (MG), endothelial cell (EC), pericyte (Per), relin-expressing Cajal-Retzius cell (CR), and two unidentified cell types (UN1; UN2). Based on the identified populations, the percentage of each cell population in the hippocampus at three different age stages and after stroke-induced injury was quantified and compared. Although some granule cells were lost in the injured hippocampus according to cell percentages, we found that pNSC and aNSC cell numbers decreased markedly with aging but were recovered in the stroke-injured hippocampus (**Figure 1E**). The average number of detected genes in each cell type is similar across different groups, thereby ruling out the possibility that the enrichment of stem cell genes is an artifact of increased global gene expression (**Figure1-figure supplement 1F)**. Overall, cell compositions and proportions varied substantially in neonatal, adult, aging and injured human hippocampus (**Figure 1E**).

**Figure 1.**
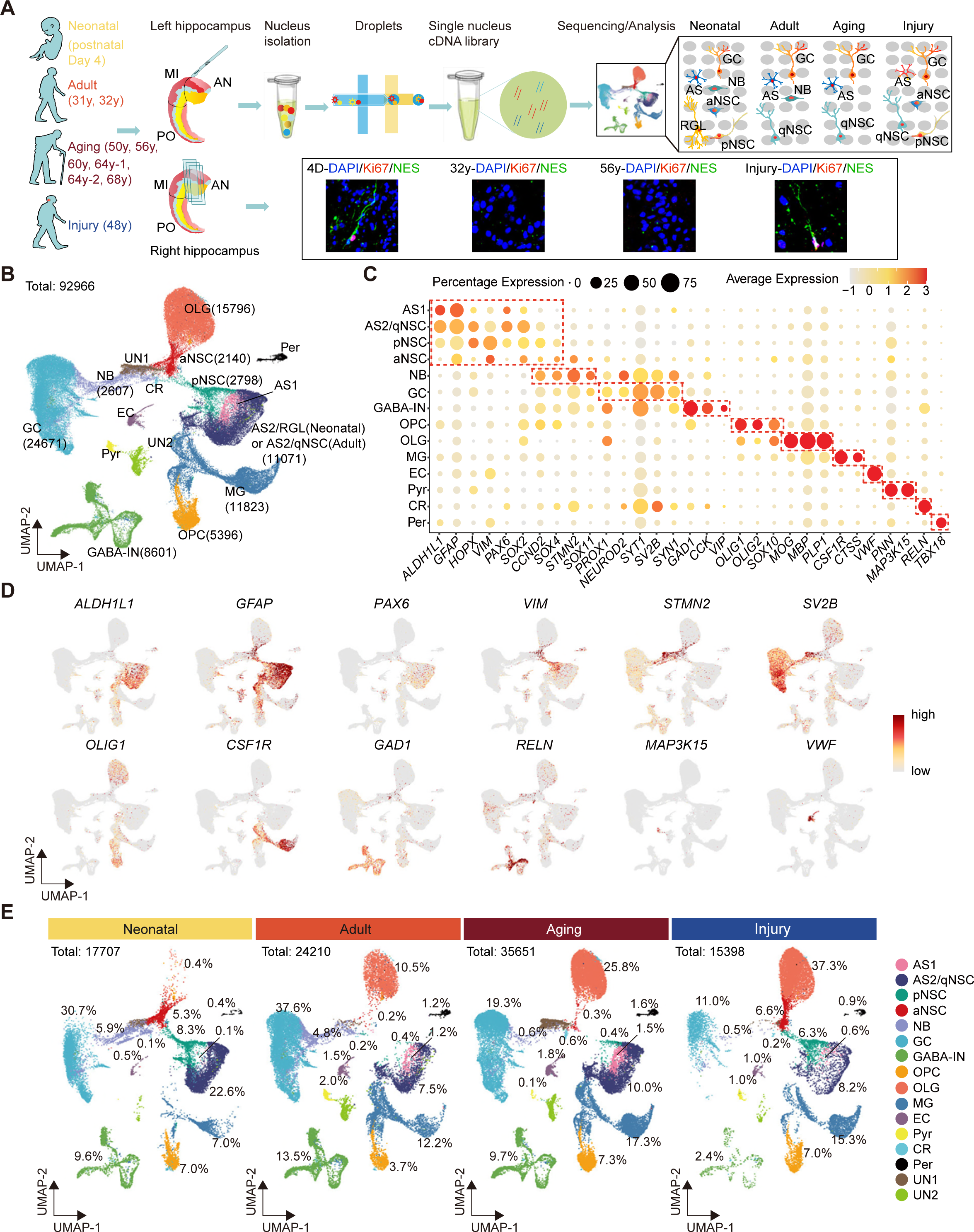
Single-nucleus transcriptomic atlas of the human hippocampus across different ages and after stroke injury. (A) Summary of the experimental strategy. The pair of hippocampi from postmortem human donors at different ages were collected. The anterior (AN) and middle (MI) parts containing dentate gyrus were used for snRNA-sequencing and immunostaining. (B) 92,966 hippocampal single nuclei were visualized by UMAP plot and categorized into 16 major populations: astrocyte1 (AS1, 1146 nuclei), astrocyte2/quiescent neural stem cell (AS2/qNSC, 11071 nuclei), primed NSC (pNSC, 2798 nuclei), active NSC (aNSC, 2140 nuclei), neuroblast (NB, 2607 nuclei), granule cell (GC, 24671 nuclei), interneuron (IN, 8601 nuclei), pyramidal neuron (PN, 676 nuclei), oligodendrocyte progenitor (OPC, 5396 nuclei), oligodendrocyte (OLG, 15796 nuclei), microglia (MG, 11823 nuclei), endothelial cell (EC, 1232 nuclei), pericyte (Per, 981 nuclei), Relin-expressing Cajal–Retzius cell (CR, 218 nuclei), and two unidentified populations (UN1 and UN2, 3810 nuclei). (C) Dot plots of representative genes specific for the indicated cell subtypes. The size of each dot represents the cell percentage of this population positive for the marker gene. The scale of the dot color represents the average expression level of the marker gene in this population. (D) UMAP feature plots showing expression distribution of cell type specific genes in all cell populations. Astrocyte (ALDH1L1, GFAP), neural stem cell (PAX6, VIM), neuroblast (STMN2), granule cell (SV2B), oligodendrocyte progenitor (OLIG1), microglia (CSF1R), interneuron (GAD1, RELN), relin-expressing Cajal-Retzius cell (RELN), pyramidal neuron (MAP3K15) and endothelial cell (VWF) are shown. Dots, individual cells; grey, no expression; red, relative expression (log-normalized gene expression). (E) Quantification of each cell population in the hippocampus at three different age stages and after stroke-induced injury. The following figure supplements are available for figure 1: **Figure supplement 1.** Cell atlas of human hippocampus across different ages and post stoke-induced injury, related to Figure 1

### The heterogeneity and molecular features of human hippocampal NSCs

Since hippocampal neurogenesis is controversial in the adult human brain (Zhong et al. 2020; Cipriani et al. 2018; Sorrells et al. 2018; Franjic et al. 2022; Boldrini et al. 2018; Moreno-Jimenez et al. 2019; Tobin et al. 2019; Terreros-Roncal et al. 2021) and the dramatic alteration of related cell types at different statuses was observed (**Figure 1E**), we mainly focused on the dissection of neural stem cells and neurogenic populations. We first performed a cross-species comparison of our human hippocampal neurogenic populations with the published single cell RNA-seq data from mouse, pig, rhesus macaque and human hippocampus (Hochgerner et al. 2018; Franjic et al. 2022). Neurogenic lineage populations across species were distributed at similar coordinates in the UMAP (**Figure 2A**). For example, human AS2/qNSCs and pNSCs aligned more strongly with astrocytes and radial glia-like cells (RGLs) from other species, and expressed classical RGL genes (**Figure 2A**). During embryonic development, pNSCs exhibit greatest similarity to RGLs. However, in the adult stage, pNSCs are in an intermediate state between quiescence and activation. Meanwhile, human aNSCs and NBs clustered together with other species’ neural intermediate progenitor cells (nIPCs) and neuroblasts, respectively (**Figure 2A**). In addition, neurogenic lineage markers identified in other species were also highly expressed in corresponding populations in our data (**Figure 2B**).

**Figure 2.**
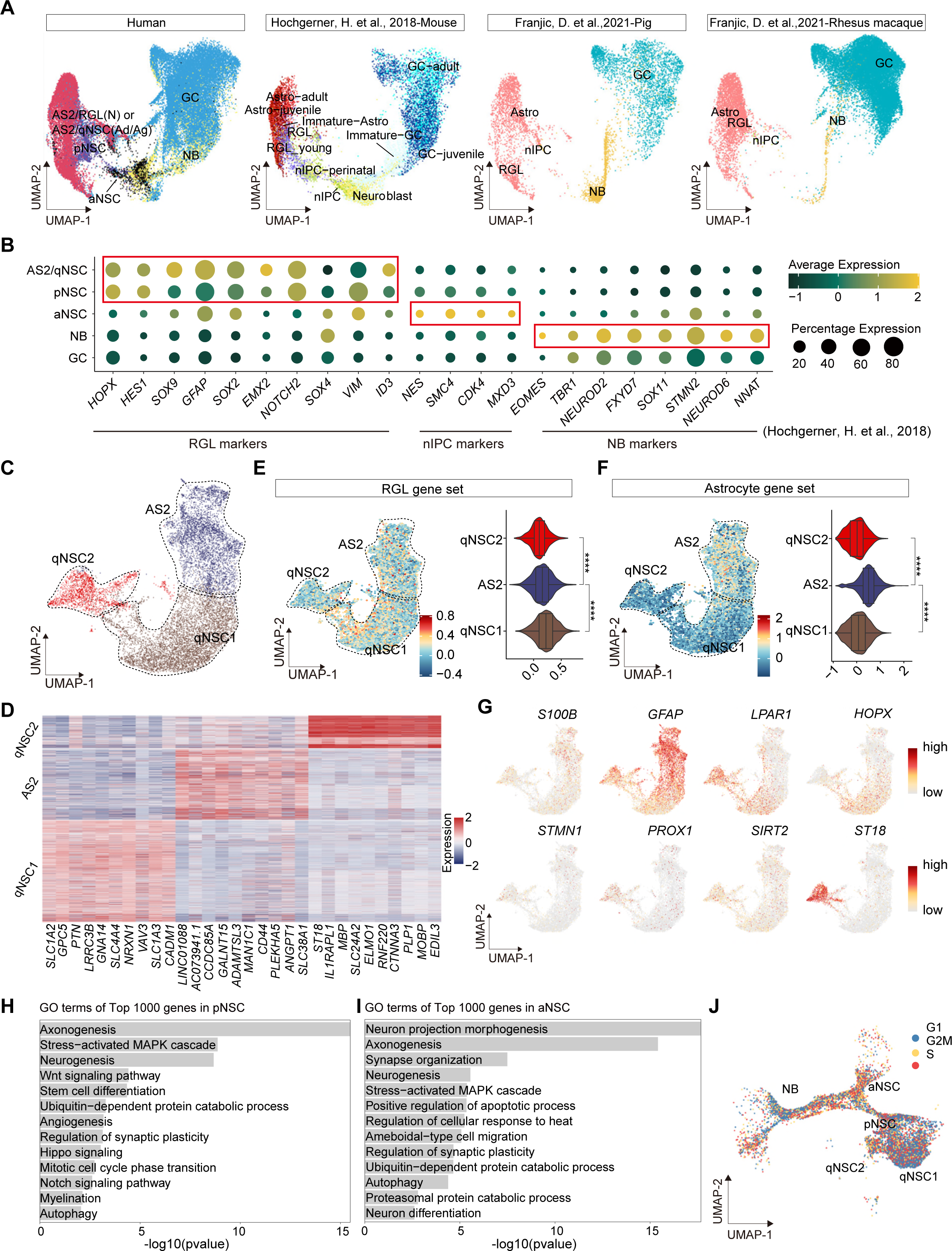
Confirmation of neurogenic lineage and dissecting of NSC molecular heterogeneity in the postnatal human hippocampus. (A) Neurogenic lineage identification was confirmed by cross-species comparison of transcriptomic signatures. Our human data were integrated with published snRNA-seq data from mice, pigs and rhesus macaque by UMAP (Hochgerner et al., 2018, Franjic et al., 2022). astrocyte2(AS2), radial glial cell (RGL), neonatal(N), quiescent neural stem cell (qNSC), adult (Ad), aging (Ag), primed neural stem cell (pNSC), active neural stem cell (aNSC), neuroblast (NB), granule cell (GC), astrocytes (Astro), neuronal intermediate progenitor cell (nIPC). (B) Expressions of previously reported RGL, nIPC, NB and immature GC markers in the corresponding populations from our human hippocampal snRNA-seq data. RGL, radial glial cell; nIPC, neural intermediate progenitor cell; NB, neuroblast; and immature GC, immature granule cell. (C) The AS2/qNSC population from neonatal sample was subclustered into three clusters, astrocyte2, qNSC1 and qNSC2. (D) Heatmap of top 10 genes (p-value < 0.05) specific for astrocytes, qNSC1 and qNSC2 after normalization. (E and F) Using Gene set scores (average, over genes in the set, of seurat function AddModuleScore) based on previously defined gene sets (Zamanian et al. 2012; Liddelow et al. 2017; Clarke et al. 2018; Hochgerner et al. 2018; Zhong et al. 2020; Franjic et al. 2022) to characterize RGL (E) and astrocytes (F). (G) UMAP feature plots showing expression distribution of cell type specific genes. Astrocyte markers (S100B and GFAP), radial glial like cell markers (*HOPX* and *LPAR1*) and neuron development markers (*ST18, STMN1, PROX1* and *SIRT2*) are shown. (H and I) Representative GO terms of the top 1000 genes specifically expressed in pNSCs (H) and aNSCs (I). (GO: BP, neural development related GO terms, p<0.05) (J) Cell-cycle phases of qNSC1, qNSC2, pNSC, aNSC and NB predicted by Cell Cycle Scoring. Each dot represents an individual cell. Steel blue, red and orange dots represent G1, S and G2/M phase cells, respectively

The following figure supplements are available for figure 2:

**Figure supplement 2.** Distinguish qNSCs and astrocytes molecular heterogeneity in the postnatal human hippocampus, related to Figure 2

qNSCs exhibit reversible cell cycle arrest and display a low rate of metabolic activity. However, they still possess a latent capacity to generate neurons and glia when they receive activation signals (Urban, Blomfield, and Guillemot 2019). They express many genes (i.e., *GFAP, ALDH1L1, VIM*) that are also expressed by astrocytes. Therefore, in our snRNA-seq data, the initial clustering (UMAP) was unable to distinguish qNSCs from astrocytes in the human hippocampus due to their high transcriptional similarity (**Figure 1B**). Previous studies in mice have shown that qNSCs express higher levels of Cd9 and Cd81 than astrocytes (E. Llorens-Bobadilla et al. 2015a), and some genes (e.g., Sox9, Hes1, Id4, and Hopx) have been proposed as essential regulators of NSC quiescence (H.M. Zhang et al. 2012; Basak et al. 2012; Giachino and Taylor 2014; Imayoshi et al. 2010; Kawaguchi et al. 2013; F. Zhang et al. 2023; Shin et al. 2015; Berg et al. 2019). However, the molecular characteristics of human qNSCs are still not well understood. To investigate the specific features of qNSCs in the human hippocampus, it is crucial to exclude astrocytes from the analysis. To this end, we performed further subclustering of the AS2/qNSC population by using Seurat (Findallmarker) analysis (**Figure 2C and D**). According to the DEGs and the feature gene expression, three subclusters were identified and annotated as astrocyte2 (AS2), qNSC1, and qNSC2 (**Figure 2C and D**). Next, we used gene set scores analysis to confirm the properties of AS2, qNSC1 and qNSC2 according to the global gene expression level (**Figures 2E and F, Figure2-figure supplement 2A and B**). Although the RGL gene set hardly distinguishes qNSCs from astrocytes (**Figure 2E**), analysis of astrocyte feature genes (Zamanian et al. 2012; Liddelow et al. 2017; Clarke et al. 2018; Hochgerner et al. 2018; Zhong et al. 2020; Franjic et al. 2022) revealed that the AS2 cluster obtained higher astrocyte score than qNSC1 and qNSC2 (**Figure 2F**). The classical astrocyte markers such as *S100B* and *GFAP* were highly expressed in the AS2 cluster (**Figure 2G**). The qNSC1 cluster was characterized by the preferred expression of quiescence NSC gene *HOPX*. Compared with the qNSC1 cluster, the qNSC2 cluster behaved less quiescent since they highly expressed *LPAR1*, neurogenic genes (e.g., *STMN1*, *PROX1*, *SIRT2* and *ST18*), showing the initial potential of lineage development (**Figure 2D and G**). It is unexpected that we observed high expression of a few OL (oligodendrocyte) genes in cluster qNSC2. However, when we compared the transcriptional similarity of qNSC2 to other populations, we still found a high correlation coefficient between qNSC2 and NSC and astrocyte populations (**Figure2-figure supplement 2C)**. We also observed that the ratio of NSCs in the astroglial lineage clusters remains higher compared to traditional histology studies. However, our data indicate a reduction in qNSCs and an increase in astrocytes during aging and injury, which supports that cell type identification by using gene set score analysis is effective, although still not optimal. Combined methods to accurately distinguish between qNSCs and astrocytes are required in the future. Compared with astrocyte and quiescent NSCs, pNSCs lowly expressed *ALDH1L1* and *GFAP*, but highly expressed stem cell genes *HOPX*, *VIM*, *SOX2*, *SOX4* and *CCND2* (**Figure 1C and D**). Consistent with their identities, gene ontology (GO) terms of the top 1000 genes in pNSCs included stem cell differentiation, Wnt signaling, neurogenesis, Notch signaling and hippo signaling, indicating that they maintain critical properties of RGLs (**Figure 2H**). Different from pNSCs, the identified aNSCs highly expressed stem cell and proliferation markers, such as *SOX2, SOX4, SOX11* and *CCND2* (**Figure 1C and D**) and were enriched for GO terms associated with the onset of neuronal fate, such as neuron differentiation, neuron projection morphogenesis, axonogenesis and synapse organization (**Figure 2I**). When we compare the differentially expressed genes between pNSC and aNSC (**Figure2-figure supplement 2D, Figure 2-source data 2**), we also found that pNSC is more associated with the Wnt signaling pathway, axonogenesis, and Hippo signaling, while aNSC is more associated with G2/M transition of mitotic cell cycle, neuron projection development, axon development, and dendritic spine organization (**Figure2-figure supplement 2E, Figure 2-source data 2**). Thus, the pNSCs referred to in this study represent an intermediate state between quiescence and activation. Different from neural stem cells, neuroblasts highly expressed *CCND2*, *SOX4*, *STMN2*, *SOX11*, *PROX1* and *NEUROD2*, and started to express several granule cell markers, such as *SYT1* and *SV2B* (**Figure 1C and D**). As expected, qNSC1 and qNSC2 were mainly in the non-cycling G0/G1 phase whereas aNSCs were mainly in the S/G2/M phase of active mitosis, confirming their quiescent and active cell states, respectively (**Figure 2J**).

Taken together, our findings demonstrate the molecular features of various types of human hippocampal NSCs and their progeny, including qNSCs, pNSCs (RGLs), aNSCs, and NBs, highlighting the heterogeneity of these cell populations and their unique cell cycle properties.

### Novel markers distinguishing various types of NSCs and NBs in the human hippocampus

The lack of validated cell-type specific markers constrains efforts to identify NSCs and their progeny in the human hippocampus. Since single-cell Hierarchical Poisson Factorization (scHPF) (Levitin et al. 2019) could sort out specific genes and seurat analysis (FindAllMarkers) is suitable for searching highly expressed genes, we used the two methods together to narrow the scope of candidate genes, allowing us to identify specific genes that can distinguish qNSCs, pNSC (RGL), aNSC and neuroblast cells from other non-neurogenic cells in the human hippocampus. The combined results from scHPF and FindAllMarkers data showed that *LRRC3B, RHOJ, SLC4A4, GLI3* were specifically expressed in qNSC1 and qNSC2, *CHI3L1* and *EGFR* could be regarded as pNSC marker genes, and *NRGN, NRN1* and *HECW1* as NB marker genes at the transcriptional level (**Figure 3A to C, Figure 3-source data 2**). Feature plots revealed that *EGFR* was specific for pNSCs, while *CHI3L1* was also expressed by astrocytes. *NRGN* and *NRN1* but not *HECW1* were specific for NBs (**Figure 3C**). Notably, several genes enriched in NBs, such as *HECW1, STMN2*, *NSG2*, *SNAP25* and *BASP1* (**Figure 3C**) were also widely distributed in GABAergic interneurons that were validated by high expression of known GABAergic interneuron marker genes, such as *DLX1, GABRG3, CCK, SLC6A11, SLC6A1, GAD1, GAD2, CNR1, GRM1, RELN* and *VIP* (**Figure 3C and D**). When we compared the granule cell lineage and the interneuron population at the whole transcriptome level between our dataset and published mouse (Hochgerner et al. 2018), macaque and human (Franjic et al. 2022) transcriptome datasets, we found high transcriptomic congruence across different datasets (**Figure 3-figure supplement 3A**). Specifically, our identified human GABA-INs very highly resembled the well-annotated interneurons in different species (similarity scores > 0.95) (**Figure 3-figure supplement 3A**). Based on the validated cell type annotation, we plotted expression of the NB/im-GC highly expressed genes reported by the other studies (Zhou et al. 2022; Wang et al. 2022b) in our identified granule cell lineage and interneuron population (**Figure 3D**). Indeed, both the previously reported genes that are regarded as the markers of NB/im-GCs, such as *DCX, PROX1,* and *CALB2* (Zhou et al. 2022; Wang et al. 2022b), and here identified genes (*HECW1, STMN2, NSG2, SNAP25* and *BASP1*) were also enriched in neonatal GABAergic interneurons (**Figure 3C and D**). Consistently, these genes were also prominently expressed in the adult interneurons (**Figure 3-figure supplement 3B**). To confirm the protein expression of DCX in interneurons, we conducted co-immunostainings of DCX and a typical interneuron marker (SST). Our results demonstrate that SST-positive interneurons are indeed capable of being stained by the traditional neuroblast marker DCX in primates (**Figure 3-figure supplement 4A-C**). These results suggest that identification of newborn neurons using NB/im-GC genes requires the exclusion of the interneuron contamination, as reported by a recent study (Franjic et al. 2022).

**Figure 3.**
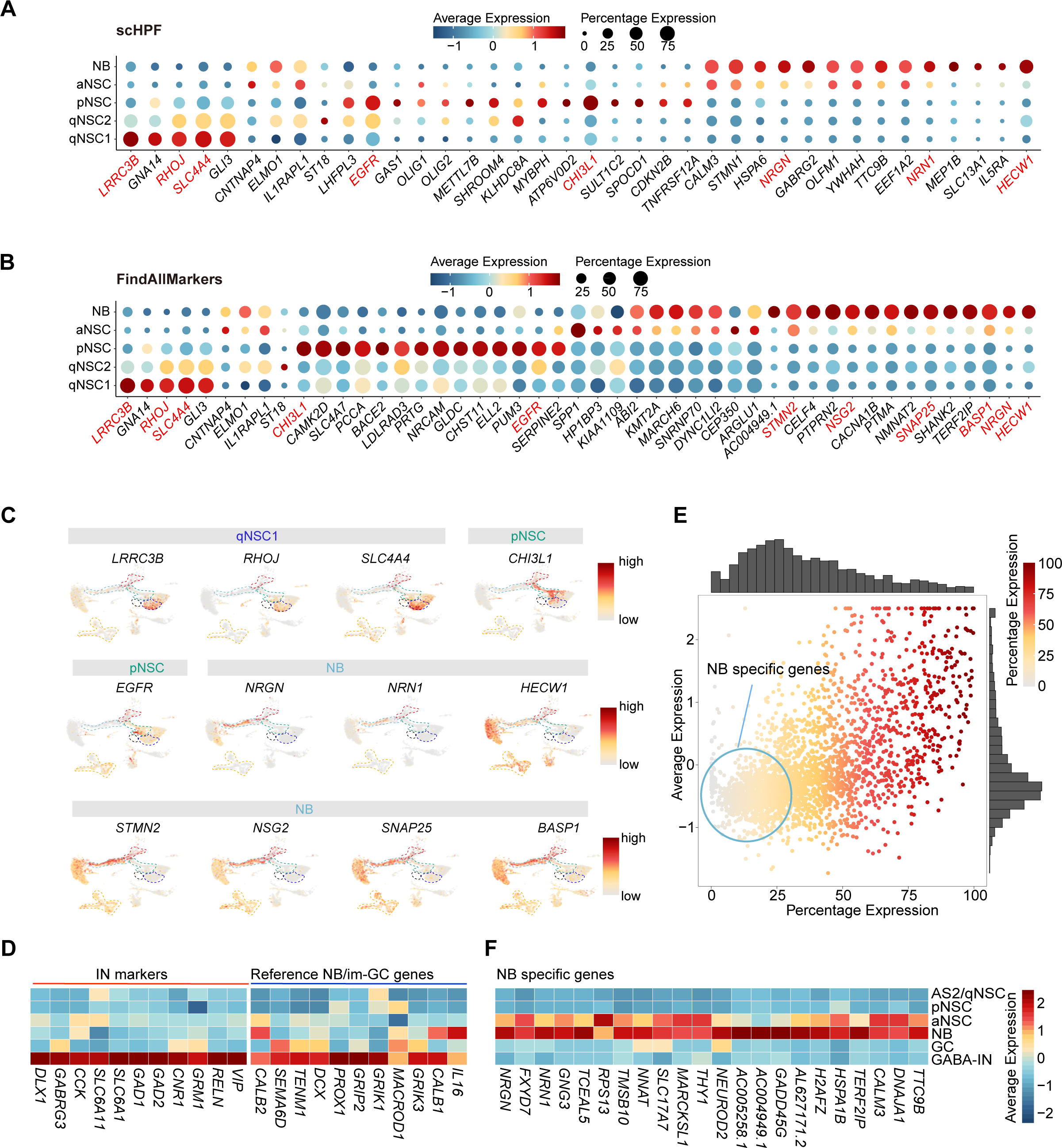
Discovery of novel markers distinguishing various types of NSCs and NBs in the human hippocampus. (A and B) Representative top genes specific for qNSC1, qNSC2, pNSC, aNSC and NB in the neonatal neurogenic lineage identified by single-cell hierarchical Poisson factorization (scHPF) (A) and FindAllMarkers function of seurat (B). (C) UMAP visualization of several cell type specific genes of the qNSCs, pNSC and NB predicted by scHPF and FindAllMarkers. (D) Heatmap showing that neuroblast/immature GC highly expressed genes that are previously reported by other literature were widely expressed in human hippocampal interneurons. (E) Scatter plot showing that several NB genes predicted by scHPF and findmarker from our snRNA-seq data were also widely expressed in human hippocampal interneurons. The genes without/with low expression in the interneurons were selected as NB specific markers (red circle scope). (F) NB specific genes selected from our snRNA-seq data were not or very low expressed in AS2/qNSCs, pNSC, GC and interneurons. The following figure supplements are available for figure 3: **Figure supplement 3.** Reported neuroblast genes were widely distributed in the adult human interneurons, related to Figure 3 **Figure supplement 4.** Neuroblast marker DCX were expressed in interneuron in macaque hippocampus of 3 months, related to Figure 3.

To further identify NB/im-GC specific genes absent in interneurons, we mapped the NB/im-GC genes identified by scHPF (top 100) and FindAllMarkers (p-adjust < 0.01) onto the interneuron population (**Figure 3E**). We selected genes with low or absent expression in the interneuron population (around the coordinate origin) as NB/im-GC specific genes by filtering out genes with high and wide distributions in the interneuron population (**Figure 3E**). We identified several representative NB/im-GC specific genes, such as *CALM3, TTC9B, NRGN, FXYD7, NRN1, GNG3, TCEAL5, TMSB10* and *NEUROD2* (**Figure 3F**) and confirmed their specificity in adult and aging samples (**Figure 3-figure supplyment 3C**).

Overall, our results revealed that most NB/im-GC genes are prominently expressed in human hippocampal interneurons, hence, our newly identified NB marker genes could be used to identify newborn neurons in adult or aging human hippocampus.

### The developmental trajectory and molecular cascades of NSCs in neonatal human hippocampus

Based on studies in mice, RGLs acquire quiescence gradually throughout postnatal development and adulthood, and share molecular markers with astrocytes (Berg et al. 2019; Alvarez-Buylla and Garcia-Verdugo 2002; Hochgerner et al. 2018; Ponti et al. 2013; Seri et al. 2001; Steiner et al. 2004; Garcia et al. 2004). The situation of RGLs in human hippocampus is still unclear. We used RNA-velocity to investigate the developmental potentials of NSCs in the neonatal human hippocampus (**Figure 4A**). We observed that pNSCs give two developmental directions: one is entering quiescence or generating AS2, and the other is generating aNSCs. Based on the GO term analysis of the differentially expressed genes comparing qNSC1/2 with pNSCs, it appears that pNSCs are more active than qNSCs (**Figure 4B**). Since qNSCs originate from RGLs (**Figure 4A**) but exit out of the cell cycle and development, the pNSC (RGL) population was set as the root of the developmental trajectory to recapitulate the continuum of the neurogenesis process (**Figure 4C**). According to the developmental trajectory, pNSCs were followed by aNSCs and NBs, and gave rise to two types of neurons (N1 and N2), indicative of ongoing neurogenesis (**Figure 4C and Figure 4-figure supplement 5A and B**). The N1 and N2 populations had distinct gene expression profiles, which indicates they are subtypes of GCs (**Figure 4-figure supplement 5C**). The N1 specifically express *NCKAP5, SGCZ, DCC, FAM19A2,* whereas the N2 specifically express *FLRT2, RIMS2, NKAIN2* and *XKR4* (**Figure 4-figure supplement 5D**).

**Figure 4.**
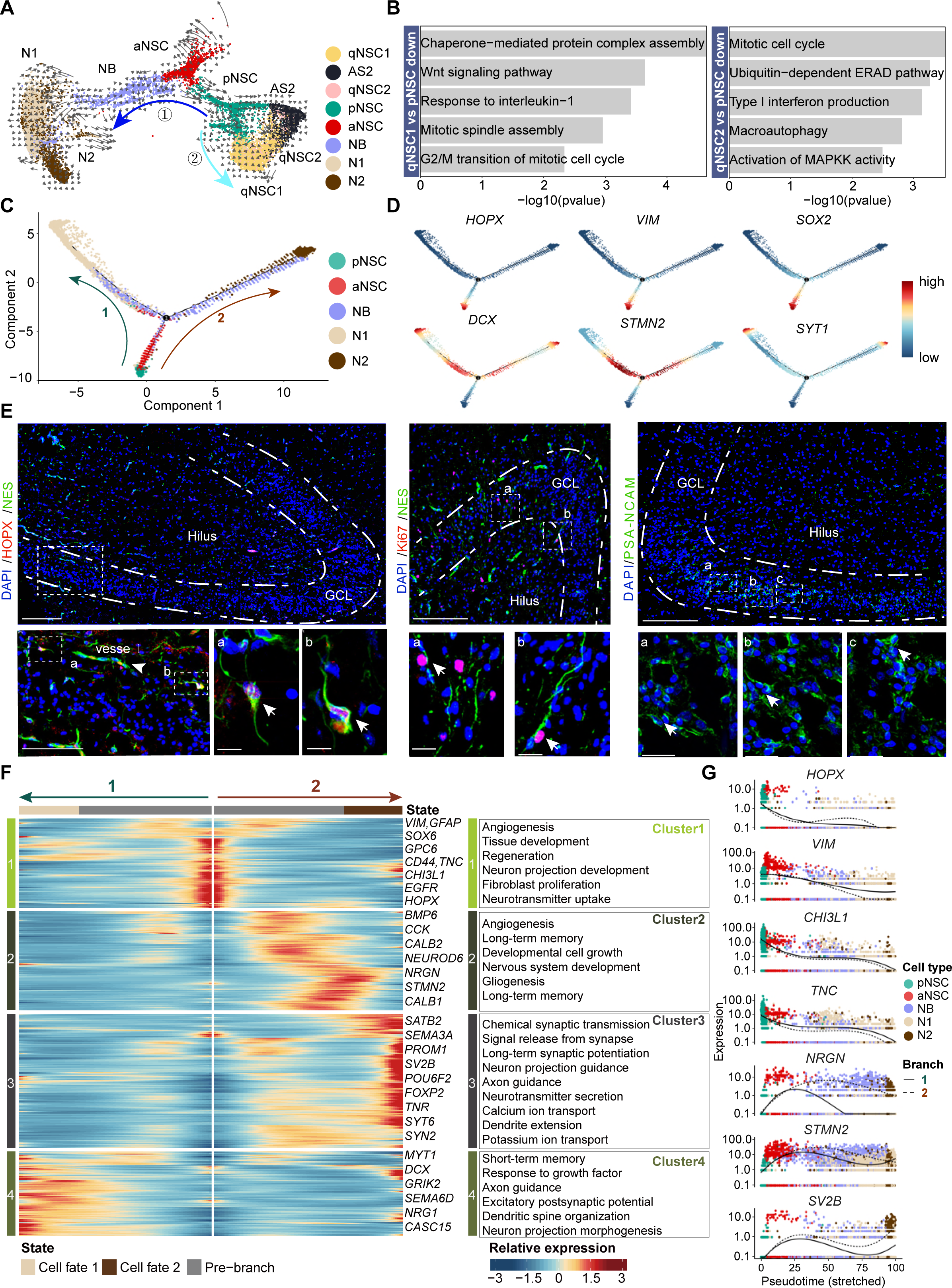
The transcriptional dynamics predicated by RNA velocity and pseudotime reconstruction revealed developmental potentials of NSC in the neonatal human hippocampus. (A) RNA velocity analysis indicating the developmental trajectory of hippocampal neurogenic lineage at postnatal Day 4. Cell types are labeled. (B) Representative GO terms of the differentially expressed genes compare qNSC1, qNSC2 with pNSC. (C) Pseudotime reconstruction of the neurogenic lineages in the neonatal human hippocampus. Dots showing individual cells. Different color represents different cell types. The arrows indicate the directions of differentiation trajectories. pNSCs as the development root was successively followed by aNSCs and neuroblasts, and then separated into two branches (1 and 2), generating two types of neuronal cells N1 and N2, respectively. (D) Expression dynamics of cell type specific genes along with the pseudotime. Each dot represents an individual cell. NSC genes (*HOPX, VIM* and *SOX2*), granule neuroblast genes (*DCX* and *STMN2*), and mature granule cell gene (*SYT1*) are shown. (E) Immunostainings of radial glia (NSC) markers (HOPX and NES), active NSC markers (NES and Ki67) and neuroblast marker (PSA-NCAM). The HOPX+NES+ RGL cells and NES+Ki67+ active NSCs with long apical processes were detected in postnatal Day4 hippocampal dentate gyrus (arrows). The PSA-NCAM+ neuroblasts (green) were located across the GCL. Scale bars of HOPX/NES immunostaining are 200 μm; the magnified and further magnified cell images are 100 μm and 10 μm, respectively; the arrowhead indicates the vessel. Scale bars of KI67/NES immunostaining are 100 μm and 10 μm, respectively. Scale bars of PSA-NCAM immunostaining are 100 μm and 10 μm, respectively; arrows indicate the neuroblasts. (F) Heatmap showing that differentially expressed genes along the pseudotime were divided into four clusters. Representative genes and enriched GO terms of each cluster are shown (GO: BP, neural development related GO terms, p<0.05). (G) Representative NSC genes (*HOPX, VIM, CHI3L1* and *TNC*), and neuronal genes (*NRGN, STMN2* and *SV2B*) were ordered by Monocle analysis along with the pseudo-time. Cell types along with the developmental trajectory were labeled by different colors. The following figure supplements are available for figure 4: **Figure supplement 5.** Pseudotime reconstruction of the neurogenic lineage development in the neonatal Day 4 human hippocampus, related to Figure 4 **Figure supplement 6.** Differentially expressed genes along the pseudotime of neurogenic lineage in the neonatal human hippocampus, related to Figure 4

Next, we traced the dynamics of pNSC, aNSC, NB and GC marker genes along with the developmental trajectory (**Figure 4D**). We found that *HOPX*, *SOX2* and *VIM* expression was preferentially maintained in pNSCs and aNSCs, but decreased upon differentiation. Finally, expressions of NB genes *DCX* and *STMN2* increased along the trajectory and the GC gene *SYT1* reached maximum expression at the end of the trajectory (**Figure 4D**). To validate the RNA-seq results, we performed immunostaining in the dentate gyrus of D4 neonatal hippocampus. In the granule cell layer (GCL) and hilus, we detected HOPX^+^NES^+^ RGLs and Ki67^+^NES^+^ proliferating NSCs (**Figure 4E**), consistent with previously reported NSC immunostaining in human hippocampus (Sorrells et al. 2018). We also found that PSA-NCAM^+^ neuroblasts located in the GCL as clusters (**Figure 4E**). These results confirm that both pNSCs (RGLs) and aNSCs maintain their proliferative status and can generate new granule cells in the neonatal human hippocampus.

To understand how gene expression profiles in different cell populations change over the developmental trajectory, we constructed gene expression cascades of neurogenesis-related cell populations (including pNSC, aNSC, NB, and N1/N2) and annotated the DEGs into 4 Clusters (**Figure 4F**). Cluster 1 population, located at the trajectory start, consists of pNSCs and aNSCs with high expression of *VIM, GFAP, SOX6, GPC6, CD44, CHI3L1, TNC, EGFR* and *HOPX.* These genes are mainly related to cell proliferation, regeneration, angiogenesis and the canonical Wnt signaling pathway (**Figure 4F, Figure 4-source data 3, Figure 4-figure supplement 6**). As expected, *HOPX, VIM, CHI3L1* and *TNC* were down-regulated along the presudotime (**Figure 4G**). Cluster 2 population highly expressed neurogenesis and neuronal differentiation genes, including *NRGN, STMN2* and *NEUROD6*, which reached their expression peak at the middle of the trajectory and represented neuron development (**Figures 4F and G, Figure 4-source data 3**). Cluster 3 and 4 populations, respectively located at the end of two branches, contained neurons that became mature and functional. The genes for the branch 2 (cluster 3) were associated with axon guidance, neurotransmitter secretion, long-term synaptic potentiation and ion transport, such as *SV2B*, *SYT6* and *SYN2*; similarly, the branch1 (cluster 4) genes were associated with neuron projection guidance, dendritic spine organization and excitatory postsynaptic potential, such as *MYT1* and *GIRK2* (**Figure 4F**). In addition, we also identified transcriptions factors (TFs) that are differentially expressed from pNSC to neurons along with the neurogenesis trajectory in all 4 clusters (**Figure 4-figure supplement 5E**. For example, progenitor cell regulation TFs *PBX3, PROX1, GLIS3, RFX4* and *TEAD1* were dominantly enriched in the origin of the trajectory. Conversely, differentiation related TFs *POU6F2, FOXP2, THRB, ETV1, NR4A3, BCL11A, NCALD, LUZP2* and *RARB* were prominently expressed in the middle and the end of the trajectory.

Our findings collectively imply that specific types of human hippocampal NSCs remain in a quiescent state postnatally, serving as a reservoir for potential neurogenesis. Meanwhile, a considerable proportion of NSCs retain their capacity for proliferation and can produce fresh granule cells in the neonatal hippocampus of humans.

### Most NSCs become deep quiescence during aging

When we quantified the cell numbers of different types of NSCs and their progeny across neonatal (postnatal day 4), adult (the mean of 31y, 32y) and aging (the mean of 50y, 56y, 60y, 64y-1, 64y-2, 68y) groups, we noted that the ratios of qNSC1 and qNSC2 increased significantly with age, particularly the deep quiescent stem cell qNSC1. Conversely, pNSC and aNSC populations sharply declined from neonatal to adult and aging stages. Meanwhile, the numbers of neuroblasts were comparable in the neonatal and adult brain, but they were markedly reduced in the aging hippocampus (**Figure 5A and B**). The abundance of neuroblasts in the adult brain suggest that compared to rodents, immature neurons in primates are indeed retained for a longer period and possess the potential to further develop into mature neurons (Kohler et al. 2011). Although the number of these neurogenic cells (pNSCs, aNSCs, NBs) in the aged hippocampus is quite low, they still expressed neural stem cell and neuroblast marker genes, including *VIM, PAX6, SOX2, PROX1, NRGN, INPP5F,* and *TERF2IP* (**Figure 5-figure supplement 7A**), ruling out that these are contaminated by other neurogenic cell types. These results together showed that pNSCs and aNSCs are present in the neonatal hippocampus, but their numbers significantly decrease with age. This suggests that human neurogenesis experiences a rapid decline after birth. In contrast, neuroblasts have a longer maturation period until adulthood, which is consistent with previous studies (Ayhan et al. 2021; Franjic et al. 2022; Ngwenya et al. 2015; Seki 2020).

**Figure 5.**
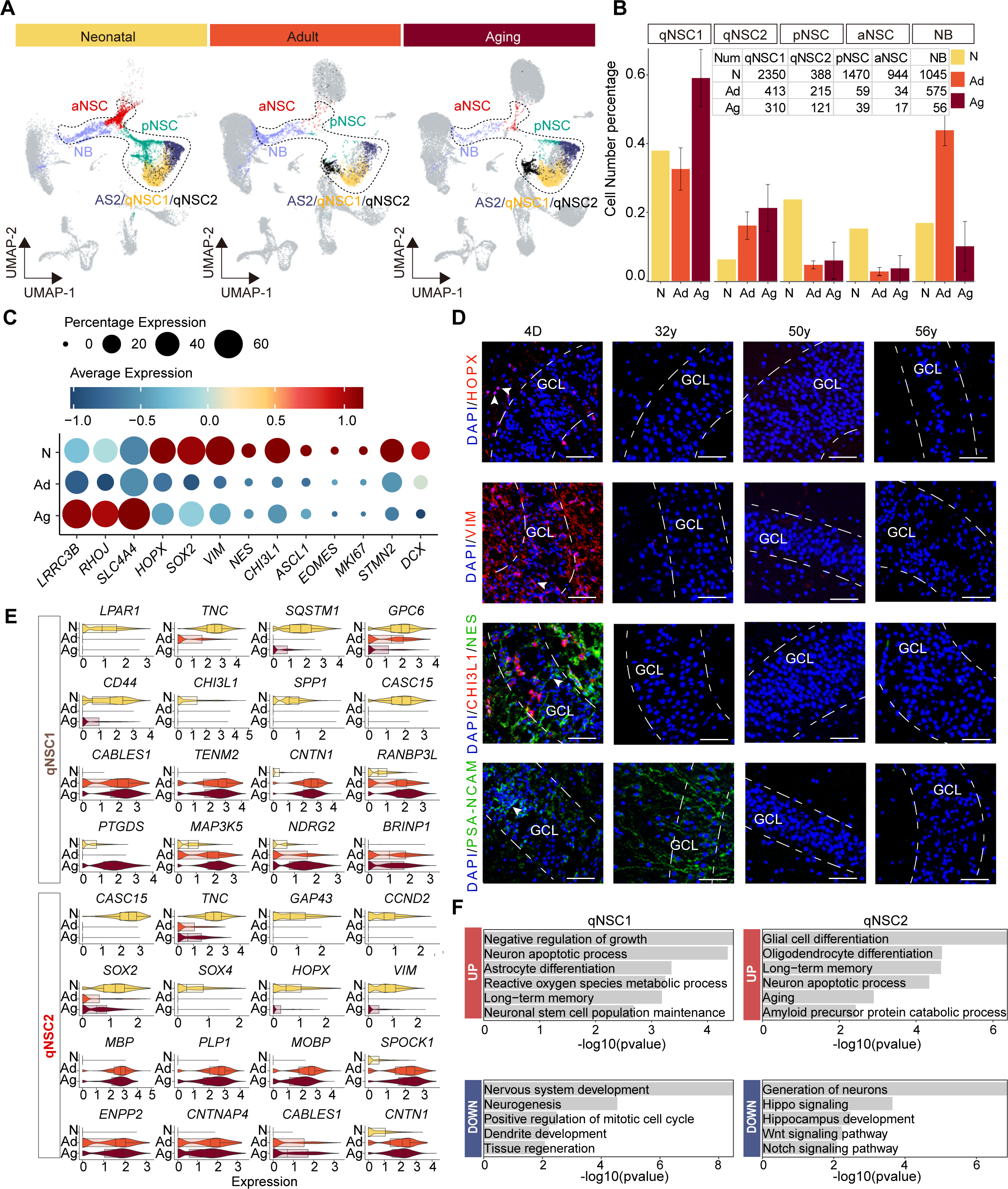
Age-dependent molecular alterations of the hippocampal NSCs and NBs. (A and B) Feature plots (A) and quantification (B) of the neurogenic populations during aging. Neonatal (N), adult (Ad), aging (Ag). The neurogenic populations include qNSC1, qNSC2, pNSC, aNSC and neuroblast. (C) The dynamic expression of some representative genes, including newly identified qNSCs genes (*LRRC3B, RHOJ,* and *SLC4A4*), NSC genes (*HOPX, SOX2, VIM, NES* and *CHI3L1*), neural progenitor or proliferation genes (*ASCL1, EOMES* and *MKI67*), and immature granule cell genes (*STMN2* and *DCX*), in human hippocampus across neonatal (D4), adult (31y, 32y) and aging (50y, 56y, 60y, 64y-1, 64y-2, 68y). (D) Immunostaining of classical NSC markers (HOPX, VIM and NES) in human hippocampal dentate gyrus across different ages (postnatal day 4, 32y, 50y, 56y). Scale bars, 60 μm. The arrowheads indicate positive cells with typical morphology. (E) Violin plot showing differentially expressed genes of qNSC1 and qNSC2 in the aging group compared to the neonatal group. (F) Representative GO terms of significantly (p-value < 0.05) up- and down-regulated genes in qNSC1 and qNSC2 during aging. The following figure supplements are available for figure 5: **Figure supplement 7.** Alterations of the neurogenic lineage related genes in human hippocampus during aging, related to Figure 5 **Figure supplement 8.** Differentially expressed genes and enrichment GO terms in pNSC, aNSC, and NB during aging, respectively, related to Figure 5

Given the continuity of NSC development and the rarity of neural stem cells in the adult or aged hippocampus, we merged the five cell types qNSC1s, qNSC2s, pNSCs, aNSCs and NBs together as the neurogenic lineage to analyze the transcriptomic alterations during aging. We have observed a significant up-regulation of astrocyte and quiescence genes (*LRRC3B, RHOJ, SLC4A4*) with increasing age, as well as a marked down-regulation of pNSC genes (*HOPX, SOX2, NES, VIM* and *CHI3L1*), aNSC genes (*ASCL1, EOMES* and *MKI67*) and NB genes (*STMN2, DCX*) upon aging (**Figure 5C**). When we stained hippocampal tissue sections from neonatal D4, 32y and 56y donors (**Figure 5D and Figure 5-figure supplement 7B**), we observed that NSC markers HOPX, VIM, NES and CHI3L1, which were widely expressed in the neonatal D4 dentate gyrus, were almost lost in 32y, 50y and 56y dentate gyrus. VIM^+^ and NES^+^ RGLs were only present around the GCL or in the hilus of the neonatal dentate gyrus (D4); whereas the NB marker PSA-NCAM was expressed in the D4 and 32y dentate gyrus, but not in 50y and 56y hippocampus (**Figure 5D**). In agreement with previous staining in adult human brain samples (Sorrells et al. 2018), PSA-NCAM^+^ cells in the GCL of the neonatal and adult dentate gyrus had neuronal morphologies (**Figure 5D**). Together, our immunostaining analysis is consistent with our snRNA-seq data, confirming that pNSCs and aNSCs experience a significant loss with aging, while neuroblasts are sustained until adulthood in humans.

To explore whether the human hippocampal NSCs are getting more and more quiescent during postnatal development and aging, we compared qNSCs from the neonatal sample with those from aged samples (**Figure 5-source data 4**). We observed that cell proliferation and growth inhibition genes (*BRINP1, CABLES1, TENM2, CNTN1*), and stem cell differentiation genes (*RANBP3L, NDRG2*) were up-regulated significantly in qNSC1 during aging. Besides *CABLES1* and *CNTN1,* the oligodendrocyte genes (*MBP, PLP1, MOBP*) were also highly expressed in aging qNSC2 (**Figure 5E**). In contrast, stem cell and regeneration genes (*LPAR1, TNC, CASC15, SOX2, SOX4, HOPX, VIM*) were down-regulated in qNSC1 and qNSC2 (**Figure 5E**). The enriched GOs of significantly up-regulated genes in aging qNSC1 and qNSC2 included negative regulation of growth, neuronal stem cell population maintenance, astrocyte differentiation, oligodendrocyte differentiation, aging and amyloid precursor protein catabolic process (**Figure 5F and Figure 5-source data 4**). Instead, the enriched GOs of significantly down-regulated genes in qNSC1 and qNSC2 were related to nervous system development, neurogenesis, positive regulation of mitotic cell cycle, tissue regeneration, autophagy, generation of neurons, hippo signaling, Wnt signaling pathway and Notch signaling pathway (**Figure 5F and Figure 5-source data 4**). All these differences between neonatal and aged qNSCs suggest that hippocampal NSCs undergo a transition into a state of deep quiescence and acquire glial properties during aging. In addition, we also compared gene expression of the remaining pNSCs, aNSCs and NBs across neonatal, adult and aged groups, respectively (**Figure 5-figure supplement 8A to I**). The DEGs and enriched GOs of each cell type also strongly revealed that neurogenesis decline with aging is mainly due to repression of neural stem cell proliferation, deficient autophagy and proteasomal protein catabolic process and increased glial cell differentiation. Overall, the results obtained from both the comparison of the entire neurogenic lineage and the comparison of individual cell types suggest that most NSCs lose their neurogenic potential as a result of entering a state of deep quiescence during aging.

### Injury induced activation of qNSCs in the adult hippocampus

The homeostasis of NSCs impacts the dynamics of neurogenesis in response to environmental signals (Chaker et al. 2015; Daynac et al. 2014; Enwere et al. 2004; Katsimpardi et al. 2014) and injury conditions in mice can even reactivate qNSCs into a proliferative state that gives rise to new neurons (E. Llorens-Bobadilla et al. 2015a; Buffo et al. 2008). In the stroke-afflicted donor (48y), we noted a significant loss of hippocampal granule neurons and interneurons (**Figure 1E**). Compared to adult donors, genes associated with apoptosis, DNA damage and autophagy were significantly up-regulated in the GCs and GABA-INs of the stroke-injured hippocampus (**Figure 6-figure supplement 9A and B**). Consistently, we detected evident cell apoptosis by TUNEL assay in the stroke-injured dentate gyrus, but not in other adult or aging samples (**Figure 6-figure supplement 9C**). These data validated that injury had occurred in the hippocampus of donors that had suffered from a stroke. Interestingly, we observed that pNSCs and aNSCs are predominantly present in the neonatal and stroke-injured samples, with minimal presence in other groups. Meanwhile, qNSCs increased with aging and reduced upon injury (**Figure 6A**). These results indicated that qNSCs may be reactivated upon injury and give rise to pNSC and aNSC populations. However, we only observed very few cells in the neuroblast population that highly expressed neuroblast marker genes *PROX1, SEMA3C, TACC2, INPP5F,* and *TERF2IP* (**Figure 5-figure supplement 7A**). We speculate that because the patient died within two days after the stroke, there was little time for the activated NSCs to generate more neuroblasts.

**Figure 6.**
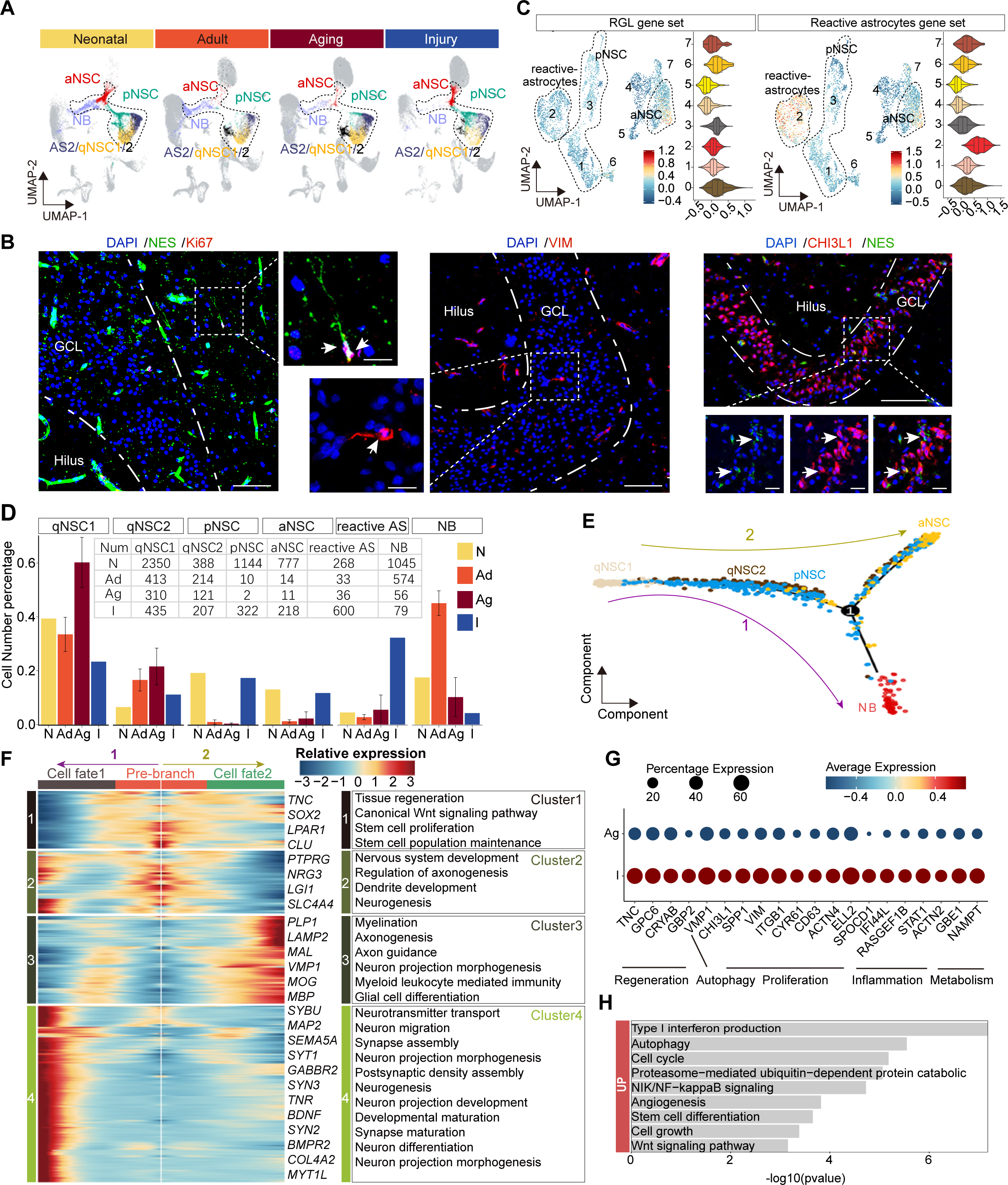
The transcriptomic signatures of the activated neurogenic lineage in the adult human injured hippocampus induced by stroke. (A) The neurogenic lineage included qNSC1, qNSC2, reactivated pNSC/aNSC and NB. Cell distribution showing by feature plots. (B) Immunofluorescence images of NES (green)/Ki67 (red), VIM (red), and CHI3L1 (red)/NES (green) showing a few active NSC cells in the 48-year-old injured hippocampal dentate gyrus. The arrows indicate radial morphology NES+/KI67+, VIM+ or CHI3L1+/NES+ active NSC cells, respectively. Scale bars, 100 μm; the magnification, 20 μm. (C) Annotated into pNSC, aNSC and reactive astrocytes according to gene set scores (average, over genes in the set, of seurat function AddModuleScore). (D) Quantification of qNSC1, qNSC2, pNSCs, aNSCs, reactive astrocytes and NB in neonatal (N), adult (Ad), aging (Ag) and stroke-injured (I) hippocampus, respectively. (E) Pseudotime reconstruction of the neurogenic lineage in the stroke-injured human hippocampus. Different colors represent different cell types. The arrow indicates the developmental direction. (F) Heatmap showing the expression profiles of differentially expressed genes (DEGs) in four clusters along the pseudotime. Representative DEGs and enriched GO terms of each cluster are shown (GO: BP, regeneration related GO terms, p<0.05). (G) The significantly up-regulated genes in neurogenic lineage upon injury compared with aging. (H) The GO term analysis of up-regulated genes in the neurogenic lineage upon injury compared with aging (GO: BP, proliferation and regeneration related GO terms, p<0.05). The following figure supplements are available for figure 6: **Figure supplement 9.** Stroke injury induced hippocampal cell apoptosis, astrocyte reactivation and neuronal damages, related to Figure 6 **Figure supplement 10.** Initially defined pNSCs and aNSCs from stroke-injured hippocampus contained reactive astrocytes and reactivated NSCs, related to Figure 6. **Figure supplement 11.** Integration of our snRNA-seq dataset with other published data, related to Figure 6.

Previous studies in mice reported that NSCs and astrocytes become activated after stroke around the injured area. Such activated NSCs which could generate newborn neurons together with reactive astrocyte-formed glial scarring may contribute to brain repair (Benner et al. 2013; Faiz et al. 2015; Li et al. 2010). Since activated NSCs and reactive astrocytes share similar transcriptional properties but have distinct morphology, we performed immunostaining of NES/KI67, NES/VIM and NES/CHI3L1 in the stroke-injured dentate gyrus. We detected a few NES^+^KI67^+^, NES^+^VIM^+^ and NES^+^CHI3L1^+^ active NSCs that had radial glia morphology with apical processes (**Figure 6B**), appearing similar to D4 NES^+^ RGLs (**Figure 4E**). However, we could not detect these active NSCs in any other adult sample (32y, 50y, 56y) (**Figure 5D**). Since VIM^+^CHI3L1^+^ reactive astrocytes with an irregular contour or star-shape morphology were widely observed in the injured hippocampus (**Figure 6-figure supplement 9E**), the pNSC and aNSC populations identified through initial UMAP clustering may include reactive astrocytes. To distinguish activated NSCs (pNSCs and aNSCs) from reactive astrocytes, we integrated neonatal pNSCs and aNSCs with injury samples, and then applied neonatal pNSC and aNSC as cell prototypes to identify pNSCs and aNSCs in the injury sample. We increased cluster resolution and obtained eight subclusters with distinct gene expression profiles (**Figure 6-figure supplement 10A and B**). When we compared the fraction of each subcluster in neonatal and injury samples, we found subclusters 0, 1 and 3 were predominant in the neonatal sample, and subclusters 2 and 4 were predominant in injury sample (**Figure 6-figure supplement 10C**). The results of Gene set score analysis also showed that subclusters 0, 1 and 3 maintained higher RGL potential than subcluster 4, and subcluster 2 had more evident reactive astrocyte properties than subclusters 0, 1 and 3 (**Figure 6C**). Consistently, RGL specific genes (*VIM*, *HOPX, LPAR1* and *SOX2*) were significantly expressed in subcluster 0, 1 and 3. The neurogenic genes (*STMN1, DCX* and *SIRT2*) were mainly expressed in subcluster 0. The reactive astrocyte marker (*OSMR, TIMP1* and *LGALS3*) were mainly expressed in subcluster 2 (**Figure 6-figure supplement 10D**). Therefore, cells in subcluster 0 were speculated as pure aNSCs, subclusters 1 and 3 were pNSC in the stroke-injured hippocampus, and cells in subcluster 2 were reactive astrocytes. Since the features of other small subclusters were not clear, they were excluded from the subsequent developmental trajectory analysis. When we quantified qNSC1, qNSC2, pNSCs, aNSCs and reactive astrocytes in neonatal (N), adult (Ad), aging (Ag) and stroke-injured (I) hippocampus, we still found that the ratios of the pNSC and aNSC populations in the neurogenic lineage reached up to 17.3% (322/1861) and 11.7% (218/1861), comparable to ratios in the neonatal group and significantly higher than the adult and aging groups. Correspondingly, the ratios of qNSC populations qNSC1 (23.4%, 413/1258) and qNSC2 (11.1%, 214/1258) in the neurogenic lineage evidently decreased in the injury group compared with adult (qNSC1=32.8%, 413/1258; qNSC2=17.0%, 214/1258) and aged group (qNSC1=57.8%, 310/536; qNSC2=22.6%, 121/536) (**Figure 6D**). These results together with the decline of neurogenesis in the aging group suggest that some quiescent NSCs in the adult and aging human hippocampus can be reactivated and give rise to active NSCs upon stroke-induced injury.

To reconstruct the injury-induced activation trajectory of qNSCs and explore their developmental potential, we excluded reactive astrocytes and included qNSC1, qNSC2, pNSC, aNSC and NB as neurogenic lineage for further analysis. In agreement with previous studies of adult neurogenesis (Dulken et al. 2017; Artegiani et al. 2017), the trajectory originated from qNSC1, qNSC2, and progressed to pNSCs, and then to aNSCs or NBs (**Figure 6E**). Based on this trajectory, *HOPX* and *PAX6* were mainly expressed where qNSCs were located, then *VIM, CD44, TNC* and *CHI3L1* reached their expression peaks in the middle of the trajectory where pNSCs located, following by *SOX2, CKAP5, RANGAP1* genes in aNSCs and *STMN2* gene in NBs (**Figure 6-figure supplement 10D**). The trajectory and gene expression together support that qNSCs can be activated to become pNSCs and aNSCs. Since the patient did not live long after the stroke, we attempted to predict the developmental potential of NSC lineages by analyzing the gene expression cascade along with the pseudotime. According to the gene expression cascade, DEGs corresponding to 4 clusters were identified (**Figure 6F**). *TNC, SOX2, LPAR1,* and *CLU* were highly expressed at the root of the trajectory (Cluster 1). Consistently, genes from the cluster 1 were related to canonical Wnt signaling pathway, tissue regeneration, stem cell proliferation and neuronal stem cell population maintenance. Subsequently, genes from Cluster 2 were enriched with the generation of neurons, dendrite and glia cell development, such as *PTPRG, NRG3, LGI1 and SLC4A4*; and lastly, genes for neuronal function (e.g., *MAP2, SEMA5A, SYT1, SYN2, SYN3 and MYT1L*) and glial fate determination (*LAMP2, PLP1, MBP* and MOG) became dominant at the end of the trajectories fate1 and fate 2 (Cluster 3 and 4) (**Figure 6F**). Accordingly, the enriched GOs of genes from Cluster 4 (fate 1) were related to neurogenesis, neuron projection development, neurotransmitter secretion, and synapse organization; the enriched GOs of genes from Cluster 3 (fate 2) were associated with glial cell differentiation and myelination (**Figure 6F and Figure 6-source data 5**). Together, our data indicate that stroke-induced injury triggers activation of qNSCs, which then generate pNSCs and aNSCs, the latter of which have the potential to give rise to either neurons or oligodendrocytes (El Waly, Cayre, and Durbec 2018; Parras et al. 2004; Enric Llorens-Bobadilla et al. 2015b; Koutsoudaki et al. 2016).

To understand relationships between regeneration and the hippocampal neurogenic lineage after stroke injury, we next explored genes involved in the activation of neurogenic lineages (qNSC1, qNSC2, pNSC, aNSC, NB). We hypothesized that genes upregulated upon injury are likely responsible for driving NSC activation. Therefore, we compared the expression of neurogenic lineage genes between the aged and injured hippocampus (**Figure 6G and H**). Specific genes that were significantly increased in the injured hippocampus were related to regeneration, autophagy, proliferation, inflammation and metabolism (**Figure 6G and H**), some of which functions have previously been demonstrated. In mice, *Tnc*, *Gpc6*, *Cryab* and *Gbp2* were reported to promote neuron regeneration and synapse formation following stroke-induced injury (Z. Chen et al. 2021; Saglam, Calof, and Wray 2021; J. Chen et al. 2010; Ugalde et al. 2020); *Vmp1*, *Chi3l1*, *Spp1*, *Vim* and *Itgb1* associated with autophagy, proliferation, and regeneration (Zhao et al. 2017; Nishimura et al. 2021; Sojan et al. 2022; Kong et al. 2018); and *Cyr61*, *CD63*, *Actn4*, *Ell2 and Spocd1* demonstrated to promote proliferation (Kong et al. 2018; Thines et al. 2022; Q. Chen et al. 2022; Alexander et al. 2017; Liu et al. 2018). Furthermore, *IFI44L, RASGEF1B* and *STAT1* are linked with both inflammation and metabolic functions (Cooles et al. 2022; Leao et al. 2020) and *ACTN2, GBE1* and *NAMPT* only for metabolic functions (Ebersole et al. 2018; Gasparrini et al. 2022). Overall, the stroke-induced up-regulated molecular signatures capture a broad activation state and regeneration of the neurogenic lineage.

## Discussion

The existence of human adult hippocampal neurogenesis has been a topic of debate over the years. Sample rarity and technical limitations are barriers that prevent us from investigating the human postnatal hippocampus during aging and post injury. With the development of snRNA-seq technology, we are able to better understand the blueprint of hippocampal neurogenesis signatures in humans. By using snRNA-seq technology, two recent studies found no adult neurogenic trajectories in human brains (Franjic et al. 2022; Ayhan et al. 2021), while other two groups newly reported that noticeable amounts of NSCs and immature neurons were found in the adult and aged human hippocampi, supporting adult human neurogenesis capacity (Zhou et al. 2022; Wang et al. 2022a). While accumulated publications support the existence of neurogenesis in the adult human hippocampus, the homeostasis and developmental potentials of neural stem cells (NSCs) under different contexts remain unclear. Here, we have revealed the heterogeneity and developmental trajectory of hippocampal NSCs, and captured its transcriptional molecular dynamics during postnatal development, aging and injury, which the traditional immunostainings could not uncover based on the limited sensitivity and specificity. Specially, we identified NSCs with different refined transcriptional statuses, including qNSC, pNSC and aNSC populations. Despite transcriptional similarity between qNSCs and astrocytes, we also distinguished qNSCs from astrocytes by using gene set score analysis.

The lack of specific markers has prevented the identification of neurogenic lineages in the human hippocampus for a long time. To fill this gap, we executed an integrated cross-species analysis combined with single-cell Hierarchical Poisson Factorization (scHPF) and seurat analysis to identify specific markers for human neurogenesis. In the study, we observed that both well-known and recently reported immature GC markers (Zhou et al. 2022; Wang et al. 2022a; Franjic et al. 2022; Hao et al. 2022), such as DCX, PROX1, and STMN2, are widely expressed in human GABAergic interneurons, which is consistent with Franjic’s observation (Franjic et al. 2022).

It suggests the risk of interneuron contamination when using these markers to identify immature GCs. We further identified new specific neuroblast markers by excluding genes expressed in human GABAergic interneurons, such as CALM3, NEUROD2, NRGN and NGN1. Thus, our findings extend our knowledge about the maker specificity of human adult hippocampal neurogenesis.

In agreement with recent studies, we also found that ETNPPL as an NSC marker (Wang et al. 2022a) was highly expressed in our identified qNSC1/2 (**Figure 6-figure supplement 11A**), and neuroblasts with the positivity of STMN1/2(Wang et al. 2022a) were maintained in the adult hippocampus (**Figure 3B and C**). In contrast, we did not find a comparable number of pNSCs, aNSCs and imGCs as reported in the aged group, but detected reactivated NSCs in the injured hippocampus. To explore the cause of the discrepancies, we examined the published human specimens’ information from different studies which reported the existence of neuroblasts in the aged hippocampi (Zhou et al. 2022). When we integrated Zhou’s snRNA-seq dataset of 14 aged donors (from 60-92 years old) with our snRNA-seq dataset, we did not detect evident pNSC, aNSC or NB populations in their 14 aged donors (**Figure 6-figure supplement 11B**). To rule out the possibility of missing cell clusters caused by analysis of Zhou’s data, we examined the expression of pNSC/aNSC markers (e.g., VIM, TNC) and neuroblast markers (e.g., STMN1 and NRGN), and they were not enriched in putative pNSC/aNSC/NB clusters, neither in other clusters (**Figure 6-figure supplement 11C and D**). However, EdU^+^PROX1^+^ newborn granule cells were observed in surgically resected young and adult human hippocampi from patients diagnosed with epilepsy, temporal lobe lesions or suspected low-grade glioma after *in vitro* culture (Zhou et al. 2022). One possibility is that these newborn granule cells were originated from the injury-induced activated NSCs caused by the process of hippocampus sectioning or *in vitro* culture. In addition, we noticed that two aged donors diagnosed with rectal cancer (M67Y) and uterine tumor (F52Y) in Wang’s study still maintained neurogenesis(Wang et al. 2022a). Given recent evidence of crosstalk between cancer and neurogenesis(Silverman et al. 2021; Mauffrey et al. 2019), we suggest that cancer might provoke neurogenesis-like status in the adult human brain. Besides, Terreros-Roncal’s work showed that amyotrophic lateral sclerosis, Huntington and Parkinson’s disease could increase hippocampal neurogenesis (Terreros-Roncal et al. 2021). Taking these data together, adult hippocampal neurogenesis is more easily to be detected in cases with neurological diseases, cancer and injuries (Terreros-Roncal et al. 2021; Zhou et al. 2022; Wang et al. 2022a). Therefore, the discrepancies among studies might be caused by health state differences across hippocampi, which subsequently lead to different degrees of hippocampal neurogenesis.

We constructed a developmental trajectory of NSCs in the neonatal hippocampus. Based on the trajectory and immunostaining analysis, we first deciphered transcriptional cascades of neurogenic lineages along with human hippocampal neurogenesis, and identified feature genes and transcriptional factors for each cell type. Combining the analysis of NSC properties and dynamics in neonatal, adult, aging and injured human hippocampus, our results supported the process of NSCs from active to quiescent status during aging and their re-activation under injury. In our study, we detected neuroblasts in the adult human hippocampus and active radial glial-like stem cells in the injured hippocampus both by immunostaining and snRNA-seq. The existence of neuroblasts but not aNSCs in the adult hippocampus indicated a long maturation period of neuroblasts in humans, in agreement with previous reports that the maturation period of neuroblasts is longer in primates than in rodents (Ngwenya et al. 2015; Seki 2020). Although a very rare number of NBs were captured by snRNA-seq, their presence was not validated by immunostaining. Because the donor died two days after the stroke, we surmise that there was not sufficient time for injury-induced aNSCs to fully differentiate into neuroblasts. However, the obviously upregulated neuronal and glial genes in the active NSC lineage in the injured hippocampus imply that these cells have the potential to generate neurons and glial cells. In addition to analyzing our own data, we also downloaded snRNA-seq data from Zhou et al. (Nature 2022), Wang et al. (Cell Research, 2022a), Franjic et al. (Neuron 2022), and Ayhan et al. (Neuron 2021) for integrative analysis. While the dataset from Zhou et al. utilized machine learning and made it difficult to extract cell type information for fitting with our own data, the datasets from the other three laboratories were successfully mapped onto our dataset. Based on the mapping analysis, AS2, qNSC, aNSC, and NB populations were identified with varying correlations in different datasets (**Figure 6-figure supplement 11 E to G**). Combined our findings and the integrative analysis, the results together suggest that the reserved qNSCs in the adult human brain can be activated by stimuli such as injury or disease, and that their inherent neurogenesis capacity can be re-awakened by specific hippocampal microenvironments.

Taken together, our work deciphers the molecular heterogeneity and dynamics of human hippocampal NSCs under different contexts. This research provides valuable insights into the development, quiescence and re-activation of human hippocampal NSCs, which may explain why adult hippocampal neurogenesis is generally difficult to observe in humans but can be detected in specific cases. However, we must acknowledge that the information about patients’ health status and relevant lifestyle parameters is limited, and the number of patients in neonatal and stroke cases is very low (n = 1). As a result, working with the current facts requires critical thinking and caution. We also realized that snRNA-seq has its limitations in distinguishing cells with very similar transcriptional signatures (such as qNSCs and astrocytes), and the function of the very rare number of NSCs or NBs that were captured by snRNA-seq without protein detection still needs to be further identified. Integrative analysis of epigenomic, proteomic and metabolomic features of individual hippocampal cells and non-invasive lineage tracing in human brain will be more valued in the future.

## Materials and methods

### Key Resources Table

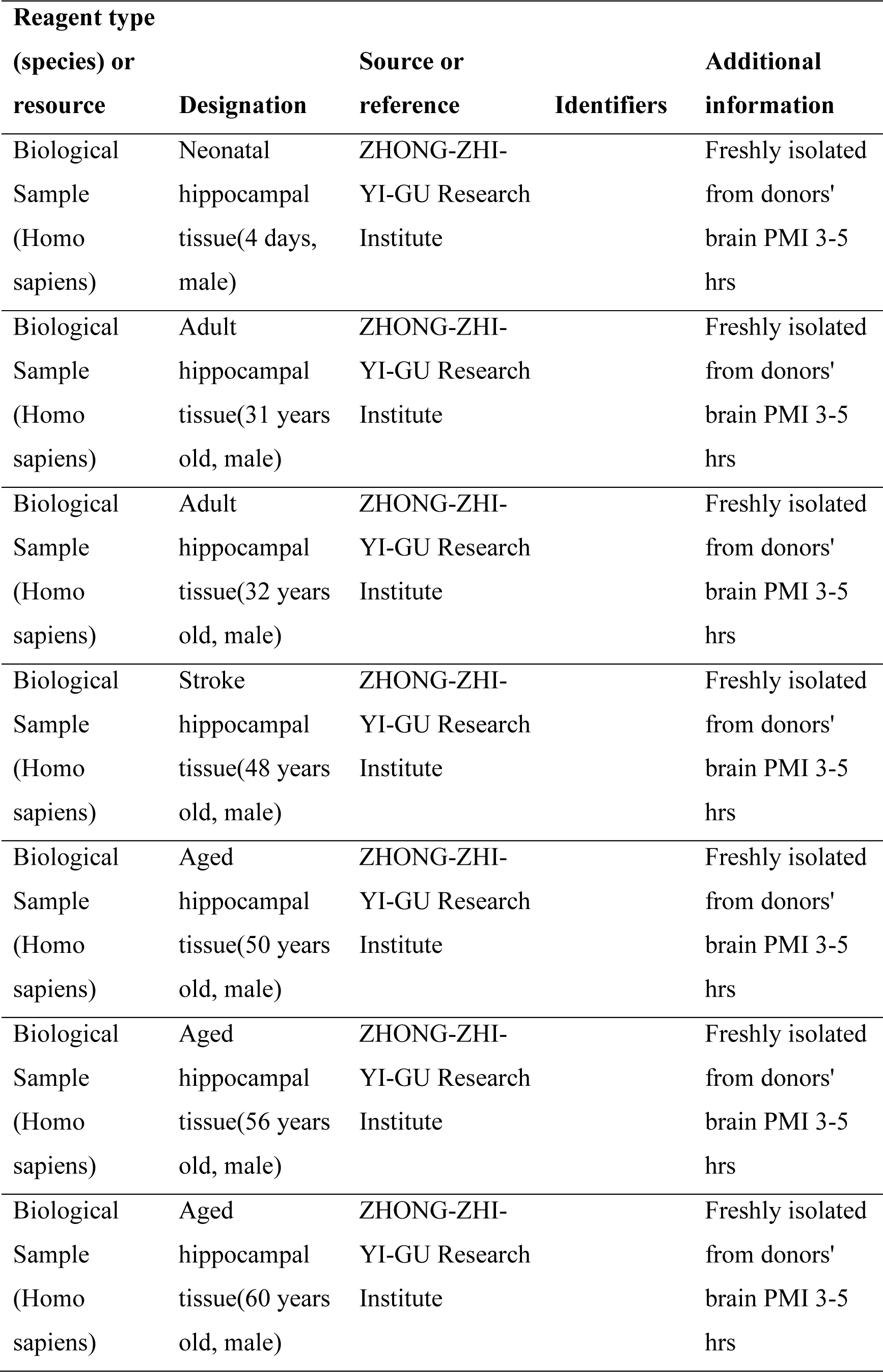
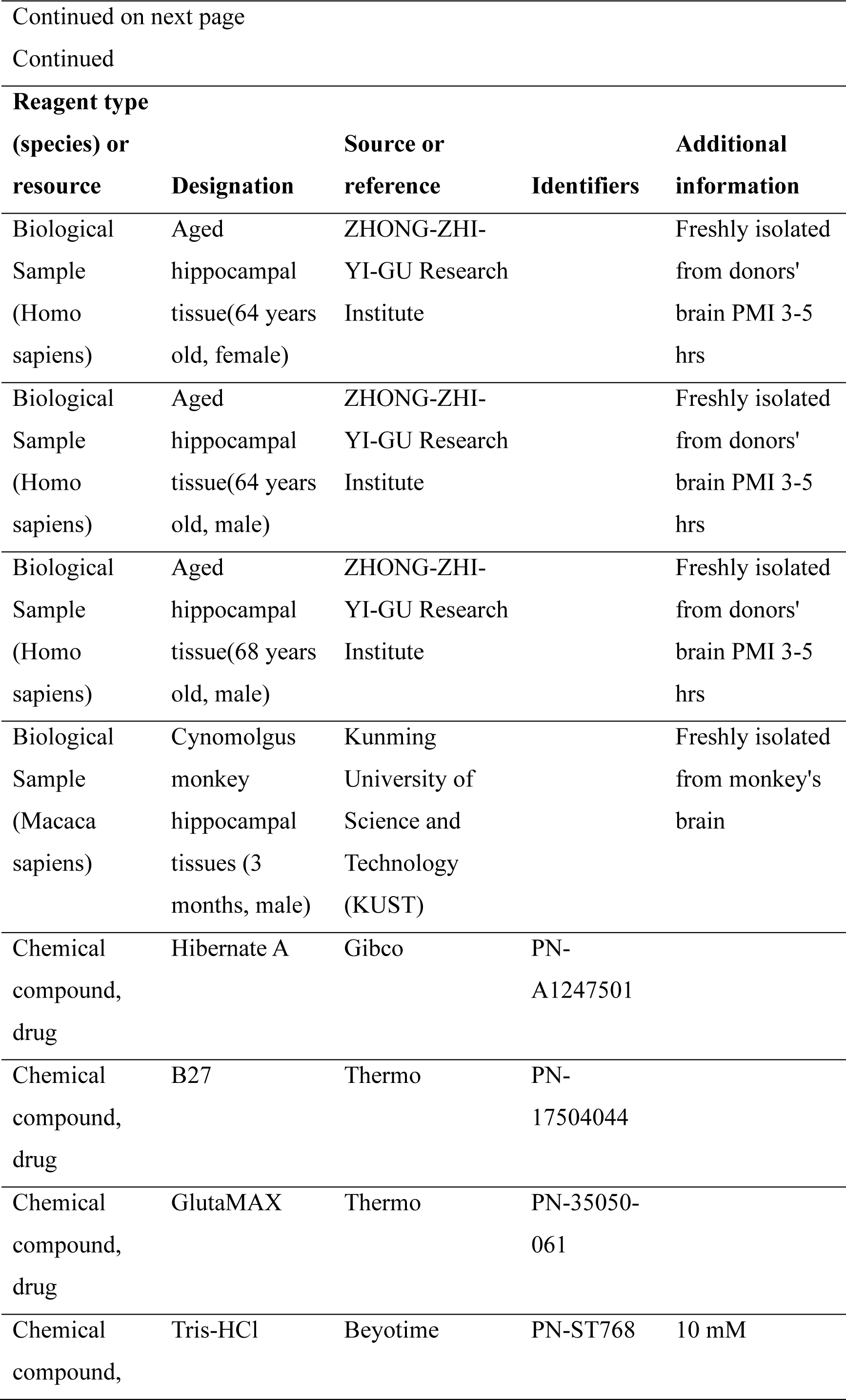
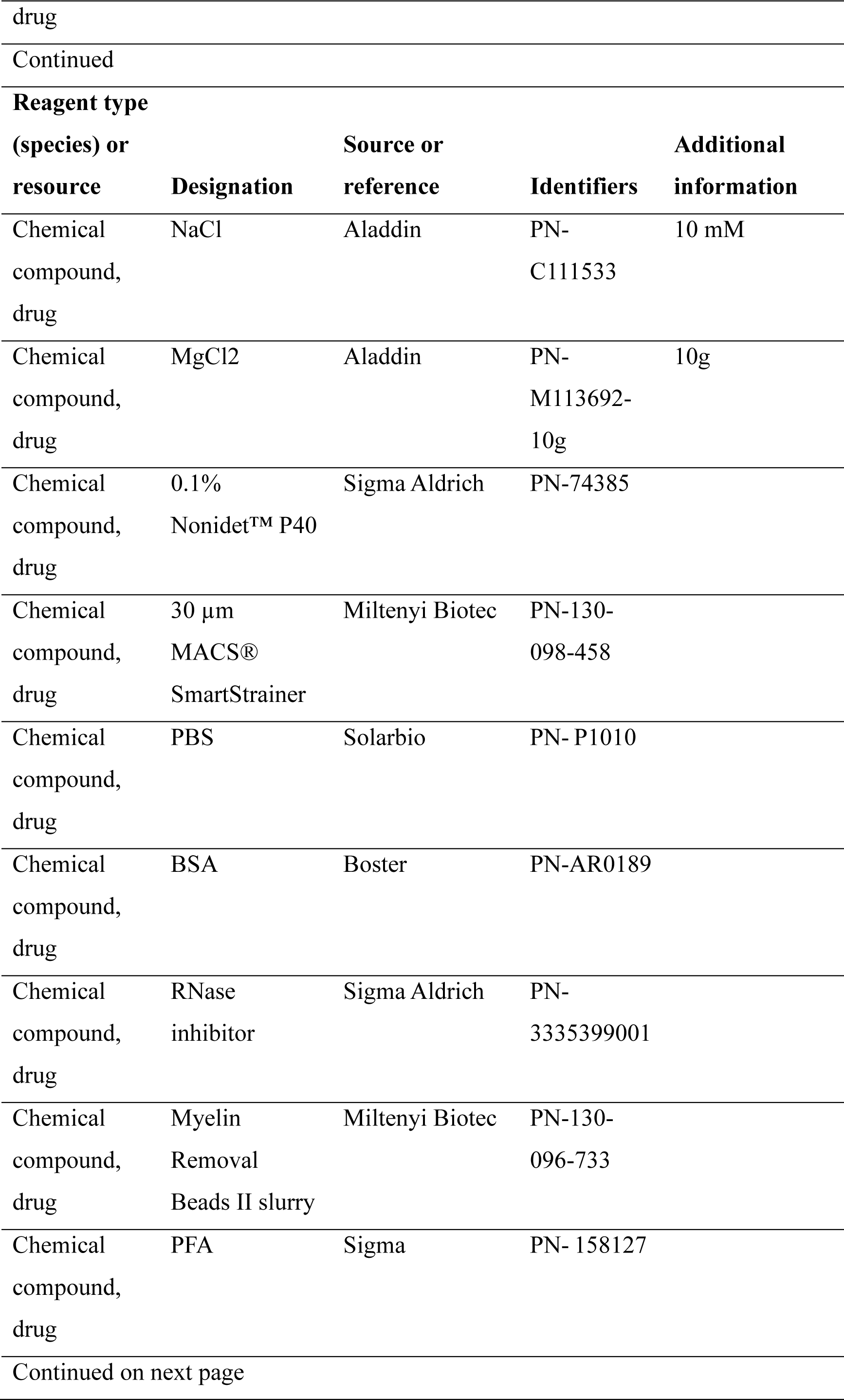
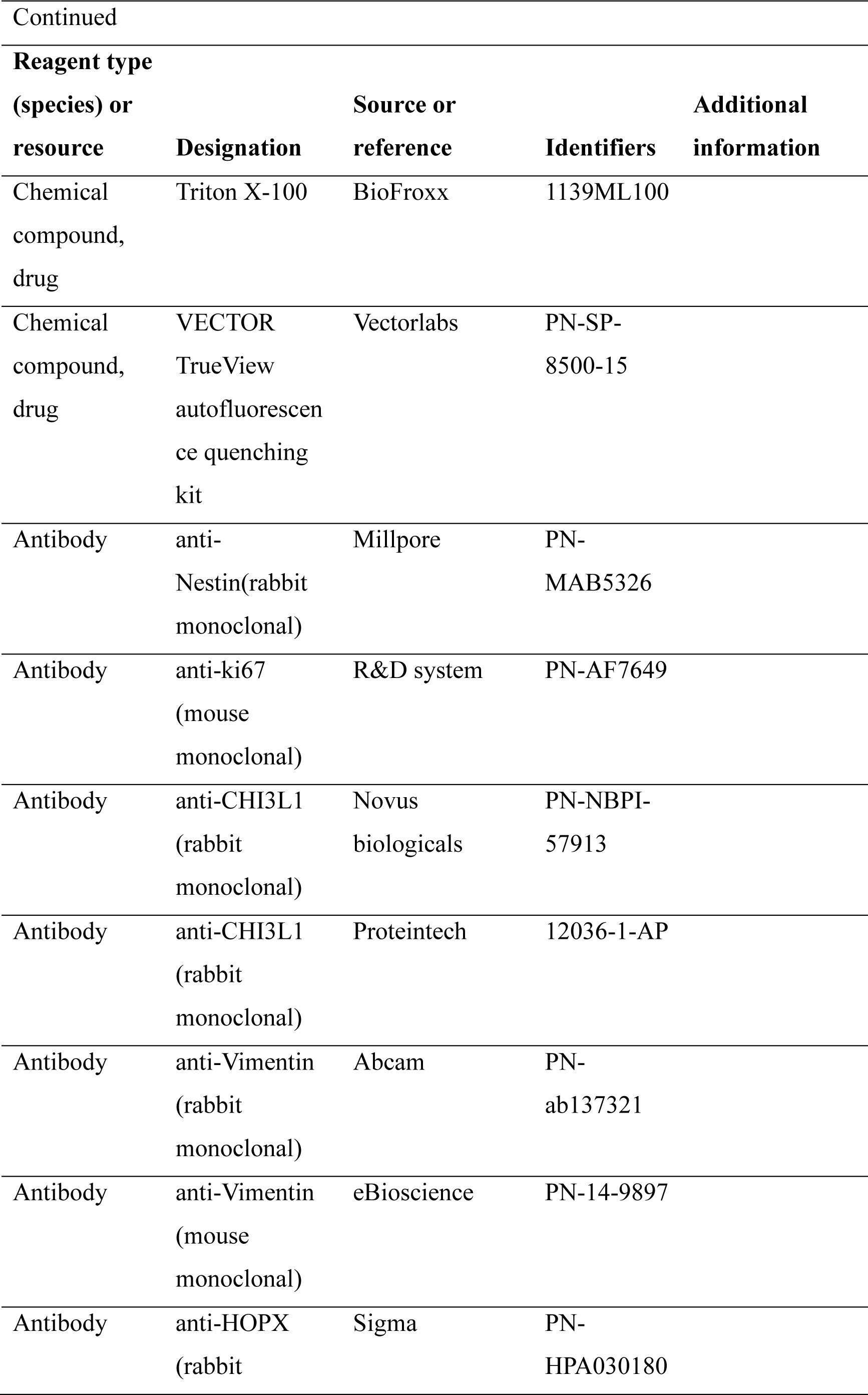
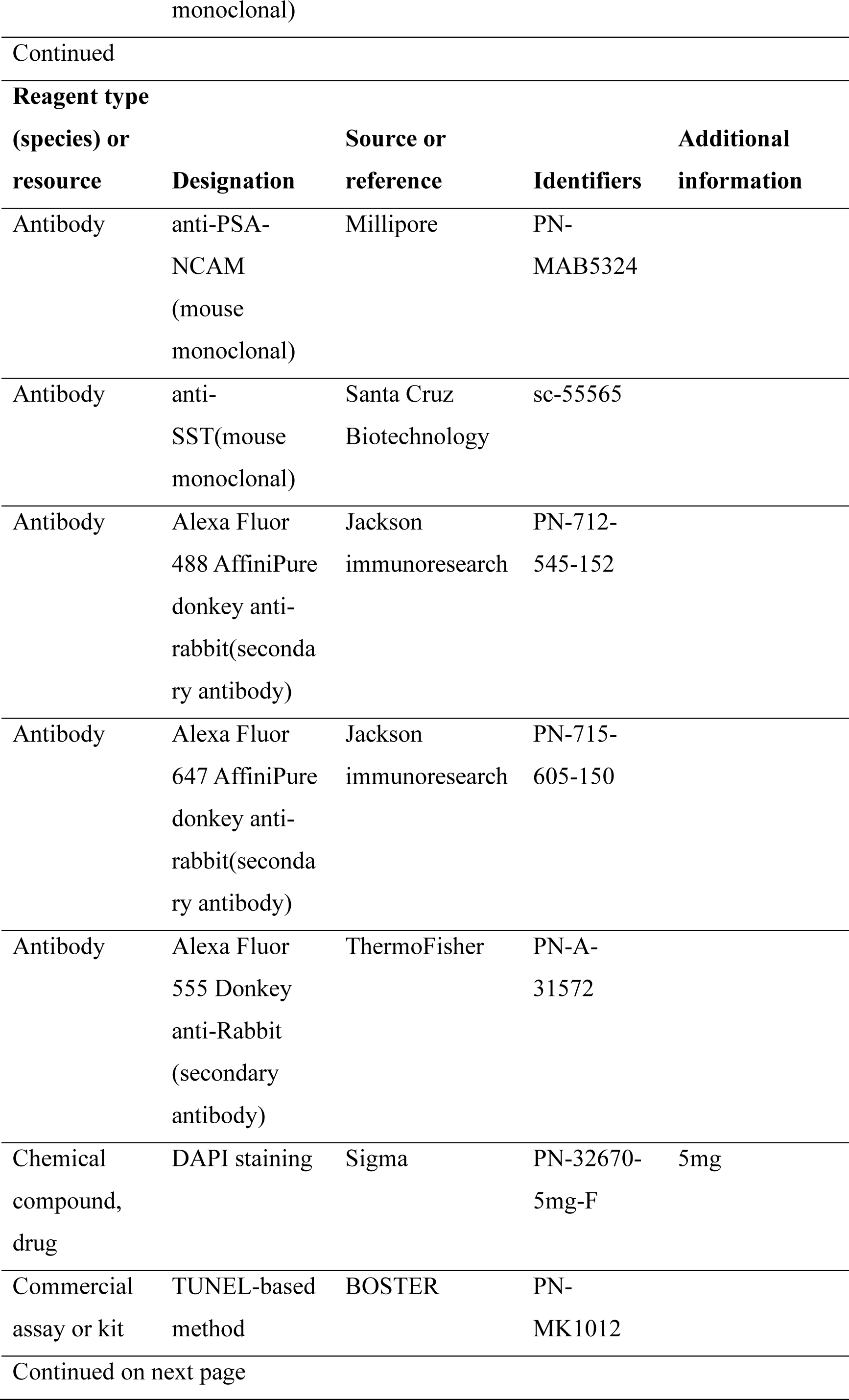
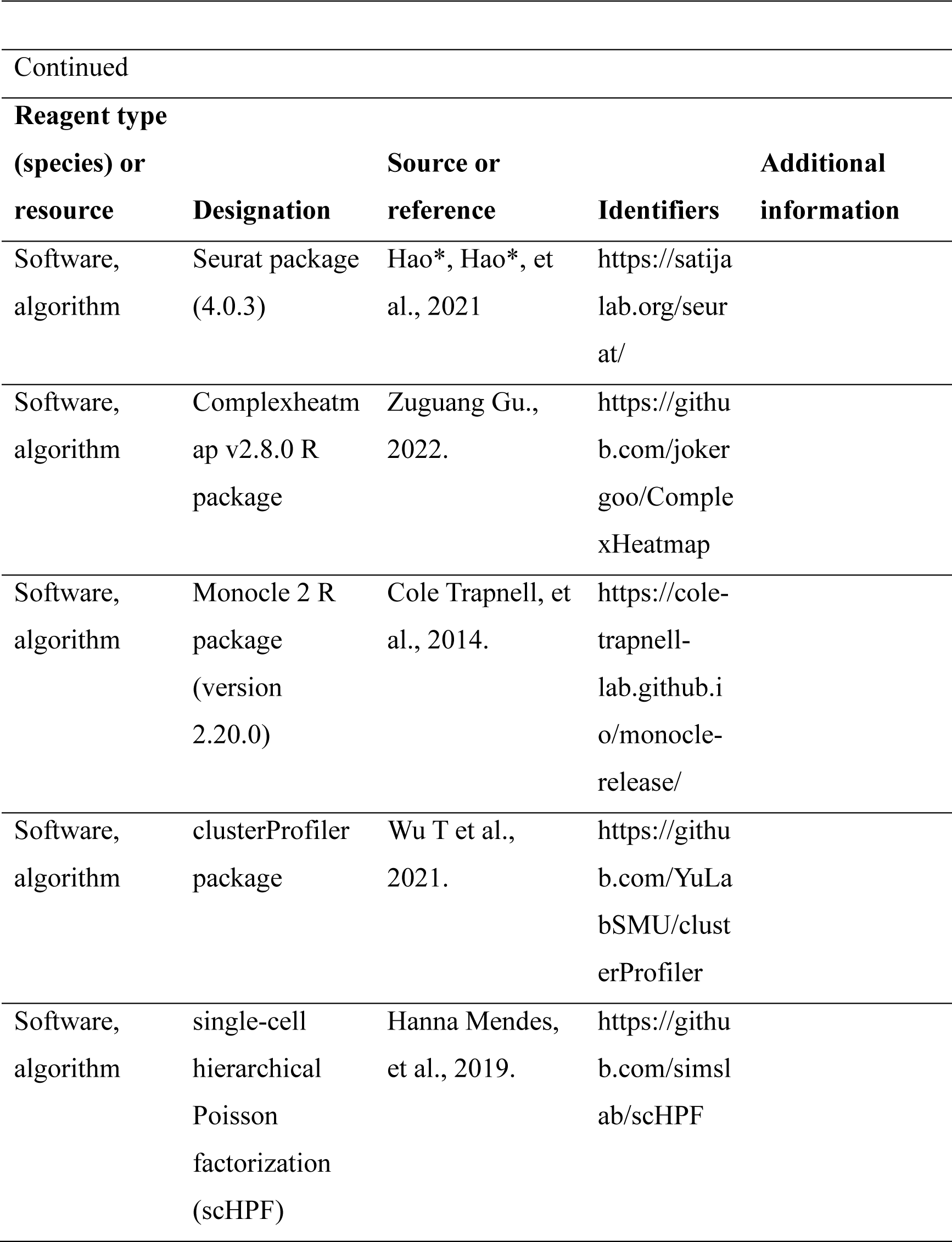

### Human hippocampal sample collection

De-identified postnatal human hippocampus samples were obtained from the ZHONG-ZHI-YI-GU Research Institute. We recruited 10 donors from neonatal day 4 to 68 years old [neonatal (postnatal 4 days), n = 1; adult (31y, 32y), n = 2; aging (50y, 56y, 60y, 64y-1, 64y-2, 68y), n = 3; stroke injury (48y), n = 1], consisting of 1 female and 9 males. Death reasons of these donors included: 1 congenital heart disease (postnatal day 4), 1 cerebral infarction (31y), 1 traumatic death (motor vehicle accident) (32y), 1 hypoxic-ischemic encephalopathy (stroke, 48y), 1 hypertension (50y), 3 carcinomas of the lungs (56y, 60y, 64y-2), 1 multiple organ failure (64y-1) and 1 carcinoma of the urinary bladder (68y) (Source Data 1). We dissected and collected the pair of hippocampi from the donors with a short post-mortem interval (about 3-4 hours). For individuals, the left hippocampus was used for sn-RNA seq analysis; the right hippocampus was fixed for immunohistochemistry analysis. Given the differences between the rostral and caudal hippocampus (Wu and Hen 2014), we used the anterior (AN) and mid (MI) hippocampus containing typical DG structures for snRNA-sequencing and immunostaining.

### Cynomolgus monkey hippocampal sample collection

Female cynomolgus monkey, in age of 3 month with body weights of 2.3 kg, was used in this study. The anterior (AN) and mid (MI) hippocampus containing typical DG structures were collected for immunostaining.

### Isolation and purification of nuclei from adult human hippocampal tissues

The cell nuclei were isolated from frozen hippocampus according to the 10x genomics nuclei isolation protocol for adult brain tissue with minor modifications (https://support.10xgenomics.com/single-cell-gene). Briefly, frozen hippocampus tissues with dentate gyrus structures were minced with surgical scissors on ice. The minced tissues were transferred into a tube with Hibernate A® (Gibco, PN-A1247501)/B27®/GlutaMAX™ (Dulken et al.) medium for equilibration. After the tissue was settled at the bottom of the tube, extra HEB was removed, leaving only enough medium to cover the tissue. Chilled lysis buffer (10 mM Tris-HCl, 10 mM NaCl, 3 mM MgCl2, and 0.1% Nonidet™ P40 Substitute (Sigma Aldrich, PN-74385) was added to the tissue and the tube was incubated on ice for 15 min with gentle shaking during the incubation. Then tissues with lysis buffer were triturated with a Pasteur pipette for 10-15 passes to obtain a single-nuclei suspension. A 30 µm MACS® SmartStrainer (Miltenyi Biotec, PN-130-098-458) was used to remove cell debris and large clumps. After centrifuging the nuclei at 500xg for 5 min at 4*°*C, Nuclei Wash and Resuspension Buffer (1X PBS with 1.0% BSA and 0.2U/µl RNase inhibitor (Sigma Aldrich, PN-3335399001) was added and gently pipetted for 8-10 times. After two times washing, Myelin Removal Beads II slurry (Miltenyi Biotec, PN-130-096-733) was added to the nuclei pellet. After resuspension and wash, the LS column and magnetic separation were applied to remove the myelin. The cleaned nuclei pellet was resuspended for density gradient centrifugation with a sucrose cushion. After centrifugation, 700-1,000 nuclei/μl was prepared for the following 10x Genomics Chromium capture and library construction protocol.

### Single nucleus RNA library preparation for high-throughput sequencing

Single nucleus RNA-seq libraries were generated by using Chromium Single Cell 3ʹ Reagent Kits v3, including three main steps: 1. GEM Generation & Barcoding; 2. Post GEM-RT Cleanup & cDNA Amplification; 3. 3ʹ Gene Expression Library Construction. Briefly, GEMs are generated by combining barcoded Single Cell 3ʹ v3 Gel Beads, a Master Mix containing cells, and Partitioning Oil onto Chromium Chip B. 5,420-18,832 nuclei were captured per channel. To achieve single nucleus resolution, nuclei were delivered at a limiting dilution. Immediately following Gel Bead-In-Emulsions (GEMs) generation, the Gel Beads were dissolved, primers containing an Illumina R1 sequence, a 16-bp 10x Barcode, a 10-bp randomer and a poly-dT primer sequence were released and mixed with cell lysate and Master Mix. After incubation of the GEMs, barcoded, full-length cDNA from poly-adenylated mRNA was generated. Barcoded, full-length cDNA was amplified via PCR to generate sufficient mass for library construction. Prior to library construction, enzymatic fragmentation and size selection were used to optimize the cDNA amplicon size. P5 primer, P7 primer, sample index sequence, and TruSeq Read 2 (read 2 primer sequence) were added via end repair, A-tailing, adaptor ligation, and PCR. The final libraries containing the P5 and P7 primers were generated by Illumina bridge amplification. Sample index sequence was incorporated as the i7 index read. TruSeq Read 1 and TruSeq Read 2 were used in paired-end sequencing (http://10xgenomics.com). Finally, the library was sequenced as 150-bp paired-end reads by using the Illumina Nova6000.

### Filtering and normalization

The Cell Ranger Single-Cell Software Suit (3.0.2) (http://10xgenomics.com) (Zheng et al. 2017) was used to perform quality control and read counting of ensemble genes with default parameters (3.0.2) by mapping to the GRCh38 pre mRNA reference genome. Only confidently mapped reads with valid barcodes and unique molecular identifiers were used to generate the gene-barcode matrix. We excluded poor quality cells after the gene-cell data matrix was generated by Cell Ranger software by using the Seurat package (4.0.3) (Butler et al. 2018; Stuart et al. 2019). Only nuclei that expressed more than 200 genes and fewer than 5000-8600 (depending on the peak of enrichment genes) genes were considered. Cells with less than 200 genes or more than 8600 genes (likely cell debris and doublets) were removed. We also removed cells with more than 20% of the transcripts generated from mitochondrial genes. The co-isolation of mitochondria during the nucleus isolation process is likely due to their association with ER. This is consistent with reports from other groups where mitochondrial DNA was detected in single-nucleus RNA-seq. In total, 33,538 genes across 92,966 single nuclei remained for subsequent analysis (Postnatal day 4 remained 17707 nuclei, 31y remained 12406 nuclei, 32y remained 11804 nuclei, 48y remained 15398 nuclei, 50y remained 5543 nuclei, 56y remained 4665 nuclei, 60y remained 7597 nuclei, 64y-1 remained 5239 nuclei, 64y-2 remained 6309 nuclei, 68y remained 6298 nuclei).

### Single-cell clustering and visualization

We used the NormalizeData and FindVariableFeatures functions implemented in Seurat V3, performed standard preprocessing (log-normalization), and identified the top 2000 variable features for each individual dataset. We then identified integration anchors using the FindIntegrationAnchors function (Satija et al. 2015).We used default parameters and dimension 20 to find anchors. We then passed these anchors to the IntegrateData function to generate integrated Seurat object. To visualize the data, we used Uniform Manifold Approximation and Projection (UMAP) to project cells in 2D and 3D space based on the aligned canonical correlation analysis. Aligned canonical correlation vectors (1:20) were used to identify clusters using a shared nearest neighborhood modularity optimization algorithm.

### Identification of cell types based on differentially expressed genes

Using graph-based clustering, we divided cells into 35 clusters using the FindClusters function in Seurat with resolution 1 (Butler et al. 2018). We identified 16 cell types including two unknown populations. The identified cell types are: astrocytes and qNSC (*GFAP, HES1, NOTCH2*), primed-quiescent neural stem cells (*HOPX, VIM*), active neural stem cells (*CCND2, SOX2*), neuroblast (*DCX, MYT1L*), granule cell (*SYT1, SV2B*), interneuron (*SST, CCK*), oligodendrocyte (*MOG*), microglia (*CSF1R*), pyramidal neurons (*PNN*), endothelial cells (*VWF*), oligodendrocyte precursor cell (*OLIG1, SOX10*), Reelin-expressing Cajal-Retzius cells (*RELN*), pericytes, and adult astrocyte (*S100B, ALDH1L1*). The DEGs of each cluster were identified using the FindAllMarkers function (thresh.use = 0.25, test.use = “wilcox”) with the Seurat R package (6). We used the Wilcoxon rank-sum test (default), and genes with average expression difference > 0.5 natural log and p < 0.05 were selected as marker genes. Enriched GO terms of marker genes were identified using enricher function with the clusterProfiler package. Hierarchical clustering and heat-map generation were performed for single cells based on log-normalized expression values of marker genes curated from literature or identified as highly differentially expressed genes. Heat maps were generated using the Heatmap function from the Complexheatmap v2.8.0 R package. To visualize the expression of individual genes, cells were grouped into different types determined by analysis with Seurat.

### Cell cycle analysis

In the cell-cycle analysis, we applied a cell-cycle related gene set with 49 genes that are higher expressed in aNSCs than in other NSCs (astrocyte-qNSC, primed NSC and neuroblast) during S and G2/M phase. UMAP plot of 92966 single-nucleus transcriptomes with points colored by putative cell-cycle phase (G0/G1, G2/M or S) using the CellCycleScoring function in Seurat (Macosko et al. 2015; Tirosh et al. 2016)

### Gene set score analysis

Gene set scores (Figures 2E and 2F) were calculated by Seurat (AddModuleScore) according to previously defined RGL cell and reactive astrocyte gene sets (Zamanian et al. 2012; Liddelow et al. 2017; Clarke et al. 2018; Hochgerner et al. 2018; Zhong et al. 2020; Franjic et al. 2022) as control feature sets. These reference raw datasets are available in the NCBI Gene Expression Omnibus (GEO) repository, accession number: GSE35338, GSE95753, GSE131258, GSE186538. Briefly, we calculated the average expression of each cell cluster on the single cell level, subtracted by the aggregated expression of control feature sets. All analyzed features are binned based on averaged expression, and the control features are randomly selected from each bin.

### Pseudo-time analysis of the neurogenic lineage in neonatal and stroke-injured hippocampal cells

The Monocle 2 R package (version 2.20.0) (Qiu, Hill, et al. 2017; Trapnell et al. 2014) were applied to construct single cell pseudo-time trajectories (Qiu, Hill, et al. 2017; Qiu, Mao, et al. 2017; Trapnell et al. 2014) to discover developmental transitions. Cells in Seurat clusters were inferred to be the corresponding locations of the neurogenesis differentiation axis. The pNSC or qNSC1 are at the beginning of pseudo-time in the first round of “order Cells”. Dispersed genes used for pseudo-time ordering were calculated by the ‘estimateDispersions’ function. “DDR Tree” was applied to reduce dimensional space and the minimum spanning tree on cells was plotted by using the visualization function “plot_cell_trajectory” for Monocle 2. (Monocle function: reduceDimension(mycds, max_components = 2, method = ‘DDR Tree’).

### Expression heatmap of highly dynamically expressed genes along the pseudotime

pNSC generated two branches, granule cell subtypes GC1 and GC2, in neonatal 4 days trajectories. These branches will be characterized by distinct gene expression programs. Branched expression analysis modeling (BEAM) aims to find all genes that differ between the branches which contain 4 gene clusters in neonatal 4 days. Differentiation-related differentially expressed genes(DEGs) were obtained with a cutoff of q value < 1 ×10^−4^, and contained 4 gene clusters. In addition, the ‘differentialGeneTest’ function in Monocle2 R package was used to find all genes that differ between trajectory cell types (qNSC1, qNSC2, pNSC, aNSC) in stroke injury hippocampus.

### Comparison of DEGs in neurogenic lineage across aging process and injury condition

We obtained significantly upregulated and down-regulated genes in aged hippocampal neurogenic lineages by comparing them with those in neonatal neurogenic lineages. Subsequently, we visualized these differentially expressed genes (DEGs) in neonatal, middle-aged and aged neurogenic lineages by violin plot and heatmap. To explore the DEGs under the stroke injury condition, we compared gene expressions of neurogenic lineages between in aged and stroke-injured hippocampus. We visualized these DEGs from neurogenic lineages in neonatal, adult, aging and stroke injury hippocampus by bubble chart to show their differential expression.

### Prediction of biological functions by GO term analysis

We enriched DEGs in neurogenic lineages during aging and under stroke injury conditions by GO term analysis. Gene ontology analysis was performed by the clusterProfiler package.

### Single cell hierarchical poisson factorization (scHPF) and seurat analysis (FindAllMarkers)

To identify new marker gene signatures associated with neurogenic lineages including qNSC1, qNSC2, pNSC, aNSC and NB in neonatal 4 days, we factorized the data with single-cell hierarchical Poisson factorization (scHPF) (Levitin et al., 2019) and seurat analysis (FindAllMarkers) from different factors onto the neurogenic lineage. To select the optimal number of factors, first, we ran scHPF for different numbers of factors, K (from 2 to 20, interval 1). Value of K 7 optimal effect. We picked the model with K=7 and presented top 10 marker genes of scHPF analysis (Figure 3A). Meanwhile, we presented top 15 gene markers of FindAllMarkers function of seurat analysis (Figure 3B).

### Multimodal reference mapping

The “Multimodal reference mapping” introduces the process of mapping query datasets to annotated references in Seurat v4. By using Seurat v4 and SeuratDisk package, we mapped Wang et al. (Cell Research, 2022a), Franjic et al. (Neuron 2022), and Ayhan et al. (Neuron 2021) human hippocampal sn-RNA datasets to our human hippocampal datasets. These reference raw datasets are available in the NCBI Gene Expression Omnibus (GEO) repository, accession number: GSE163737, GSE186538, GSE160189. First, annotate each query cell based on a set of reference-defined cell states. Second, project each query cell onto our previously computed UMAP visualization.

### Immunostaining of human and monkey hippocampal tissues

The hippocampus from the right side of the human brain with a short post-mortem interval was dissected. Monkey is deeply sedated with isoflurane and then euthanized with an overdose of pentobarbital. The monkey brain was removed from the skull, and the hippocampus was obtained. The human and monkey hippocampal tissue fixed with 4% paraformaldehyde (PFA) for up to 24 hours and cryoprotected in 30% sucrose at 4 °C until completely sink to the bottom. The tissue samples were frozen in OCT (Tissue-Tek) on dry ice and sectioned at 10 μm on a cryostat microtome (Leica CM1950). Tissue slides sectioned from the anterior of the hippocampus containing typical dentate gyrus structures were first incubated in blocking and permeation solution with 2% Triton X-100 (Sigma) for 2 h. Next, the sections were treated with a VECTOR TrueView autofluorescence quenching kit (Vectorlabs, PN-SP-8500-15) to reduce the innate auto-fluorescence of the human tissue, washed with 3×15min PBS (pH 7.6), and then incubated in 3% bovine serum albumin (BSA) for 1 hour at RT. Subsequently, sections were incubated overnight at 4℃ with the following primary antibodies: anti-Nestin (rabbit, 1:500, Millpore, PN-MAB5326) and anti-ki67 (mouse, 1:500, R&D system, PN-AF7649); anti-CHI3L1 (rabbit, 1:200, Novus biologicals, PN-NBPI-57913; rabbit, 1:100, Proteintech, 12036-1-AP); anti-Vimentin (rabbit, 1:300, Abcam, PN-ab137321); anti-Vimentin (mouse, 1:800, eBioscience, PN-14-9897); anti-HOPX (rabbit, 1:500, Sigma, PN-HPA030180); anti-PSA-NCAM (mouse, 1:500, millipore, PN-MAB5324); SST, mouse (sc-55565, Santa Cruz Biotechnology). After overnight incubation, tissue sections were washed with PBS for 3×15min, and then incubated with secondary antibodies at room temperature for 2 hours: Alexa Fluor 488 AffiniPure donkey anti-rabbit IgG(H+L) (1:500, Jackson immunoresearch, PN-712-545-152), Alexa Fluor 647 AffiniPure donkey anti-rabbit IgG(H+L) (1:500, Jackson immunoresearch, PN-715-605-150). DAPI staining (Sigma, PN-32670-5mg-F), Donkey anti-Rabbit IgG (H+L) Highly Cross-Adsorbed Secondary Antibody, Alexa Fluor™ 555(1:500, ThermoFisher, PN-A-31572) was performed and sections were washed with 1 X PBS for 3 x 15min. After washing, sections were mounted and dried, ready for microscope observation.

### Terminal deoxynucleotidyl transferase dUTP nick end labeling (TUNEL) assay

Tissue sections were analyzed for DNA fragmentation using a TUNEL-based method (BOSTER, PN-MK1012). Briefly, sections were first permeabilized in 0.02% Triton X-100 overnight. To label damaged nuclei, 20 μL of the TUNEL reaction mixture (Labeling buffer, TdT, BIO-d-UTP) was added to each sample and kept at 37 °C in a humidified chamber for 120 min. Sections were a wash with PBS for 2 min and blocked with 50 μL blocking reagent at room temperature for 30 min. Then SABC buffer and DAPI were added following the protocol of BOSTER TUNEL kit for the detection of apoptotic cells.

## Acknowledgements

This work was supported by the National Key Research and Development Program of China (2022YFA1103100), the Key Research Project of Science and Technology of Yunnan (YNWR—YLXZ—2020—015), the National Natural Science Foundation of China (NSFC) (32070864 and 32160153), STI2030-Major Projects (2022ZD0207700), Xingdian Talent Support Program (KKRD202273100), Major Basic Research Project of Science and Technology of Yunnan (202001BC070001 and 202102AA100053) and the Natural Science Foundation of Yunnan Province (202105AD160008, 202101AT070287 and 202207AA110003).

## Author contributions

T.L. conceptualized, initiated and organized the project. T.L., R. Zhang and C.Z. supervised the project. T.L., R. Zhang and J.Y. designed experiments. J.Y. and R. Zhang performed experiments and helped with bioinformatics analysis. S.D. and R. Zhu analyzed the RNA-seq data. J.T., J.S., J.X. and C.H. collected hippocampus tissue. Q.Z., X.D., L.G. and J.L. performed the tissue staining and imaging. Y.Y. and N.L. prepared a single nucleus RNA library. T.L., R. Zhang. and J.Y. wrote the manuscript.

## Human hippocampal tissues and ethics statement

This work was approved by the ZHONG-ZHI-YI-GU Research Institute of Human Research Protection (ZZYG-YC2019-003). All donated tissues in this study were from dead patients. Tissue was collected following the guidelines recommended by the Ethical Review of Biomedical Research Involving People for tissue donation. Hippocampus tissue samples were collected after the donor patients (or family members) signed an informed consent document that was in strict observance of the legal and institutional ethics at ZHONG-ZHI-YI-GU Research Institute. All hippocampal samples used in these studies had not been involved in any other procedures. All the protocols followed the Interim Measures for the Administration of Human Genetic Resources, administered by the Ministry of Science and Technology of China.

## Cynomolgus monkey hippocampal tissues and ethics statement

Animal ethics statement Female cynomolgus monkeys, in age of 3 month with body weights of 2.3 kg, were used in this study. All animals were housed at Kunming University of Science and Technology (KUST), and individually bred in an American standard cage at a light/dark cycle of 12 hours/12 hours. Reference Number of the Research Ethics Committee, Kunming University of Science and Technology: KUST202301005. All animal procedures were approved in advance by the Institutional Animal Care and Use Committee of Kunming University of Science and Technology and were performed in accordance with the Association for Assessment and Accreditation of Laboratory Animal Care International for the ethical treatment of primates.

## Data and code availability

The accession numbers for the raw snRNA-seq data reported in this paper in Genome Sequence Archive (GSA): HRA003049. Specimen information and sequencing statistics are described in Source data1. The code used to perform analyses in this paper is available on GitHub at https://github.com/BigreyR/snRNA-seq_hippocampus.

## Conflicts of interest

The authors declare that they have no competing interests.

The code used to perform analyses in this paper is available on GitHub at https://github.com/BigreyR/snRNA-seq_hippocampus.

## Source code files

**1 Figure 1-Source Code1: Single-nucleus transcriptomic atlas of the human hippocampus across different ages and after stroke injury.**

1.1 Fig1B Atlas umap

1.2 Fig1C Cell marker bubble plot

1.3 Fig1D Cell marker feature plot

1.4 Fig1E Atlas umap split by Stage

**2 Figure 1-figure supplement 1 related to Figure 1: Cell atlas of human hippocampus across different ages and post stoke-induced injury.**

2.1 FigS1AB QC Violin

2.3 FigS1C Atlas umap(3D)

2.4 FigS1D Atlas umap color by samples

2.5 FigS1E Top marker heatmap

2.6 FigS1F Mean gene summary by celltype

**3 Figure 2-Source Code2: Confirmation of neurogenic lineage and dissecting of NSC molecular heterogeneity in the postnatal human hippocampus.**

3.1 Fig2A Cross-species comparison umap

3.2 Fig2B Hochgerner_H.et_al.2018 Marker check

3.3 Fig2C AS_qNSC subtype umap

3.4 Fig2D AS_qNSC subtype top marker heatmap

3.5 Fig2EF AS_qNSC subtype Addmodulescore

3.6 Fig2G AS_qNSC subtype Featureplot

3.7 Fig2HI aNSC_pNSC top1000 GOBP

3.8 Fig2J Cell-cycle phases umap

**4 Figure 2-figure supplement 2 related to Figure 2: Distinguish qNSCs and astrocytes molecular heterogeneity in the postnatal human hippocampus.**

4.1 FigS2A AS_qNSC subtype Addmodulescore split by stage

4.2 FigS2B AS_qNSC subtype cellpercent split by stage

4.3 FigS2C qNSC2_OLG corr heatmap

4.4 FigS2D pNSC vs aNSC DEGs heatmap

4.5 FigS2E pNSC vs aNSC DEGs GOBP barplot

**5 Figure 3-Source Code3: Discovery of novel markers distinguishing various types of NSCs and NBs in the human hippocampus.**

5.1 Fig3AB find new marker by scHPF & Findallmarker

5.3 Fig3C New marker Featureplot

5.4 Fig3DF NB_GC markers heatmap

5.5 Fig3E Percent & Expression pointplot

**6 Figure 3-figure supplement 3 related to Figure 3: Reported neuroblast genes were widely distributed in the adult human interneurons.**

6.1 FigS3A corr with ref heatmap

6.2 FigS3BC NB GABA-IN specific bubble split by age

**7 Figure 3-figure supplement 4 related to Figure 3: Neuroblast marker DCX were expressed in interneuron in macaque hippocampus of 3 months.**

**8 Figure 4-Source Code4: The transcriptional dynamics predicated by RNA velocity and pseudotime reconstruction revealed developmental potentials of NSC in the neonatal human hippocampus.**

8.1 Fig4A neonatal NSCs RNA velocity

8.3 Fig4CD neonatal pseudotime trajectory & gene expression

8.4 Fig4F neonatal pseudotime branch heatmap

8.5 Fig4G neonatal pseudotime branch gene expression

**9 Figure 4-figure supplement 5 related to Figure 4: Pseudotime reconstruction of the neurogenic lineage development in the neonatal Day 4 human hippocampus.**

9.1 FigS5AB Neurogenic_lineage pseudotime trajectory

9.2 FigS5C N1 N2 specific Marker heatmap

9.3 FigS5D N1 N2 specific Marker Featureplot

9.4 FigS5E N1 N2 pseudotime branch heatmap

**10 Figure 4-figure supplement 6 related to Figure 4: Differentially expressed genes along the pseudotime of neurogenic lineage in the neonatal human hippocampus.**

10.1 FigS6ABCD Neurogenic_lineage pseudotime branch gene expression

**11 Figure 5-Source Code5: Age-dependent molecular alterations of the hippocampal NSCs and NBs.**

11.1 Fig5A NSCs_NB Age-dependent umap

11.2 Fig5B NSCs_NB Age-dependent cellpercent

11.3 Fig5C NSCs_NB Age-dependent marker bubble

11.5 Fig5E qNSC1_qNSC2 Age-dependent DEGs violinplot

11.6 Fig5F qNSC1_qNSC2 Age-dependent GOBP barplot

**12 Figure 5-figure supplement 7 related to Figure 5: Alterations of the neurogenic lineage related genes in human hippocampus during aging.**

12.1 FigS7A pNSC_aNSC_NB marker bubbleplot split by age

**13 Figure 5-figure supplement 8 related to Figure 5: Differentially expressed genes and enrichment GO terms in pNSC, aNSC, and NB during aging, respectively.**

13.1 FigS8ADG pNSCs aNSCs NB Age-dependent DEGs

13.2 FigS8BCEFHI pNSCs aNSCs NB Age-dependent GOBP

**14 Figure 6-Source Code6: The transcriptomic signatures of the activated neurogenic lineage in the adult human injured hippocampus induced by stroke**

14.1 Fig6A NSCs_NB Injury-related umap

14.2 Fig6C NSCs_NB Injury-related Addmodulescore

14.3 Fig6D NSCs_NB Injury-related cellpercent

14.4 Fig6E NSCs_NB Injury-related pseudotime trajectory

14.5 Fig6F NSCs_NB Injury-related pseudotime branch heatmap

14.6 Fig6G NSCs_NB Injury-related DEGs Bubble

14.7 Fig6H NSCs_NB Injury-related DEGs enrichment

**15 Figure 6-figure supplement 9 related to Figure 6: Stroke injury induced hippocampal cell apoptosis, astrocyte reactivation and neuronal damages.**

15.1 FigS9A GC_IN Injury-related GOBP barplot

15.2 FigS9B GC_IN Injury-related DEGs violinplot

**16 Figure 6-figure supplement 10 related to Figure 6: Initially defined pNSCs and aNSCs from stroke-injured hippocampus contained reactive astrocytes and reactivated NSCs.**

16.1 FigS10ABC Injury-related reactive subtype umap

16.2 FigS10D Injury-related reactive subtype Featureplot

16.5 FigS10E NSCs_NB Injury-related pseudotime gene expression

**17 Figure 6-figure supplement 11 related to Figure 6: Integration of our snRNA-seq dataset with other published data.**

17.1 FigS11A Check NSC_NB new marker by Wang’s data

17.1 FigS11B Integration Zhou’s refdata umap

17.1 FigS11CD Integration Zhou’s refdata featureplot

17.1 FigS11E Multimodal reference mapping wang_et_al

17.1 FigS11F Multimodal reference mapping Franjic_et_al

17.1 FigS11G Multimodal reference mapping Ayhan_et_al

## Notes

### Competing Interest Statement

The authors have declared no competing interest.

### Summary of Updates

In this revised version, we have added the limitations of this work in the last paragraph of the discussion section. We also updated the title, key resource table, funding information, and list of source code files in the main text document.

