## Supplementary figures and legends for "Deciphering molecular heterogeneity and dynamics of human hippocampal neural stem cells at different ages and injury states"

### Supplement figure legends

**Figure 1-figure supplement 1. Cell atlas of human hippocampus across different ages and post stoke-induced injury.** (A) The raw sequencing data revealed that an average of more than 2000 genes per cell were detected, with mitochondrial genes comprising less than 5%. The detailed data for each sample can be found in **source data 1**. This indicates that the hippocampal tissue used for analysis exhibits good cellular viability and high RNA integrity. (B) Excluded cells with mRNA expression of less than 200 genes or more than 8600 genes (potential cell debris and doublets). Additionally, cells with over 20% of generated transcripts from mitochondrial genes were also removed. (C) Visualization of major cell types from human hippocampal snRNA-seq data by using 3D UMAP. (D) Cell atlas of each human hippocampal sample. Different colors indicate different samples. (E) Heatmap of top 50 genes (p-value < 0.05) specific for each major population after normalization. AS1, adult astrocyte; AS2/qNSC, astrocyte/quiescent neural stem cell; pNSC, primed neural stem cell; aNSC, active neural stem cell; NB, neuroblast; GC, granule cell; GABA-IN, GABAgeric-interneuron; Pyr, pyramidal neuron; OPC, oligodendrocyte progenitor cell; OLG, oligodendrocyte; MG, microglia; EC, endothelia cell; Per, pericyte; CR, Relin-expressing Cajal-Retzius cell; Unknown1 (UN1); Unknown2 (UN2). (F) The average number of genes detected in each cell type are similar across different groups. Related to Figure 1.

**Figure2-figure supplement 2. Distinguish qNSCs and astrocytes molecular heterogeneity in the postnatal human hippocampus**

(A) Using Gene set scores (average, over genes in the set, of seurat function AddModuleScore) based on previously defined gene sets (Zamanian et al. 2012; Liddelow et al. 2017; Clarke et al. 2018; Hochgerner et al. 2018; Zhong et al. 2020; Franjic et al. 2022) to distinguish qNSCs and astrocytes from AS2/qNSC population in different groups concluding neonatal, adult, aging and injury. (B) Quantification of the respective proportions of qNSCs and astrocytes by taking the sum of astroglia lineage (qNSC1, qNSC2, AS2 and AS1) across different groups. Neonatal (N), adult (Ad), aging (Ag), injury(I). qNSC1, qNSC2, AS2 and AS1 (Adult-AS). (C) Transcriptional congruence observed between neurogenesis and gliogenesis populations is demonstrated by the matrix plot scores, which indicate the similarity between each pair of populations. (D) The differentially expressed genes of aNSC compared with pNSC. (E) The GO analysis of differentially expressed genes of aNSC compared with pNSC (GO: BP, neural development related GO terms, $p<0.05$).
Related to Figure 2.

**Figure 3-figure supplement 3. Reported neuroblast genes were widely distributed in the adult human interneurons.** (A) Transcriptional congruence of granule cell lineage and interneuron population between our dataset and published mouse, macaque and human transcriptome datasets. The matrix plot indicates the similarity scores of given human hippocampal cell populations from our dataset (rows) assigned to the corresponding literature-annotated cell types (columns). (B) Neuroblast genes reported by several literatures were widely distributed in the adult human interneurons from 10 individuals. (C) Our identified neuroblast specific genes were absent in the adult human interneurons. Related to Figure 3.

**Figure 3-figure supplement 4. The neuroblast marker DCX was expressed in interneurons ($SST^+$) in the hippocampus of 3-month-old macaques.** (A-C) Co-immunostainings of the neuroblast marker (DCX) and the interneuron marker (SST) on macaque hippocampal tissue were showed with different magnification views. Since we encountered a lack of DCX immunostaining signaling in our collected human hippocampal tissue, we opted to utilize the hippocampal tissue from 3-month-old macaques instead. In the images, some of the SST+ interneurons (green) with long apical processes were co-stained with DCX (red). Scale bars for panel A are 200 μm, while the magnified cell images in panels B and C are 50 μm. The arrowhead indicates the colocalization of DCX and SST. Related to Figure 3.

**Figure 4-figure supplement 5. Pseudotime reconstruction of the neurogenic lineage development in the neonatal Day 4 human hippocampus.** (A) Cells during neurogenic lineage development were ordered by Monocle analysis along pseudo-time. (B) Cell types located at developmental trajectory were labeled by different colors. (C and D) Heatmap (C) and UMAP (D) visualization of distinct differentially expressed genes between N1 and N2. (E) Heatmap showing expression dynamics of transcriptional factors (TFs) along the neurogenesis trajectory. The representative TFs in each cluster were shown on the right.

**Figure 4-figure supplement 6. Differentially expressed genes along the pseudotime of neurogenic lineage in the neonatal human hippocampus.**
Representative NSC genes, and neuronal genes were ordered by Monocle analysis along with the pseudo-time. Cell types along with the developmental trajectory were labeled by different colors. (A) Representative genes in Cluster 1. (B) Representative genes in Cluster 2. (C) Representative genes in Cluster 3. (D) Representative genes in Cluster 4. Related to Figure 4F.

**Figure 5-figure supplement 7. Alterations of the neurogenic lineage in human hippocampus during aging.** (A) Bubble plots showing our identified pNSCs, aNSCs and NBs from 10 individuals still express neural stem cell and neuroblast marker genes during aging despite their rare number. 48y donor was a stroke sample. (B) Immunostainings of classical NSC markers (HOPX, VIM and NES), pNSC gene CHI3L1 identified by us and neuroblast marker PSA-NCAM in human hippocampal dentate gyrus across different ages (postnatal day 4, 32y and 56y). Scale bars: 4D-500 μm, 32y-800 μm, 56y-600 μm. Related to Figure 5.

**Figure 5-figure supplement 8. Differentially expressed genes and enrichment functions in pNSC, aNSC, and NB along aging, respectively.** (A) Heatmap showing differentially expressed genes (DEGs) across neonatal, adult, aged pNSC (p-value < 0.05). (B and C) Representative GO terms of significantly up-regulated (B) and down-regulated (C) genes during pNSC aging (GO: BP, selected neural development related GO terms from Top 200, p-value < 0.05). (D) Heatmap showing DEGs across neonatal, adult, aged aNSC (p-value < 0.05). (E and F) Representative GO terms of significantly up- regulated (E) and down-regulated (F) genes during aNSC aging (GO: BP, neural development related GO terms, p-value < 0.05). (G) Heatmap showing DEGs across neonatal, adult, aged NB (p-value < 0.05). (H and I) Representative GO terms of significantly up- regulated (H) and down-regulated (I) genes during NB aging (GO: BP, neural development related GO terms, p-value < 0.05). Related to Figure 5.

**Figure 6-figure supplement 9. Stroke injury induced hippocampal cell apoptosis, astrocyte reactivation and neuronal damages.** (A) Representative GO terms of upregulated genes in stroke-injured hippocampal GCs and INs, compared with the normal aged hippocampus (GO: BP, selected stress and immune response related GO terms from Top 200, p-value < 0.05) (B) Genes relative to apoptosis, DNA damage and autophagy were significantly upregulated in the stroke-injured hippocampus, compared with the normal aged hippocampus. I, injury; Ag, aging. (C) TUNEL assay showing obvious cell apoptosis in the 48y dentate gyrus, but not in other adult samples, which confirmed the stroke caused hippocampal injury. Scale bars, 500 μm. (D) CHI3L1 and VIM co-immunostaining showing some CHI3L1+VIM+ and CHI3L1+VIM- cells exhibited morphologies of reactive astrocyte (arrowhead) and neuron (arrow) in the GCL and hilus, respectively. Scale bars, 500 μm; the magnified images, 100 μm and 20 μm. Related to Figure 6.

**Figure 6-figure supplement 10. Initially defined pNSCs and aNSCs from stroke-injured hippocampus contained reactive astrocytes and reactivated NSCs.** The integrative analysis of single cell data was based on initially defined pNSC and aNSC populations from the neonatal Day 4 and 48y-stroke injury hippocampus. (A) Integrative analysis of pNSC and aNSC from stroke injury and neonatal hippocampus showing that these cells were subclustered into 8 clusters. (B) Heatmap of top 10 genes (p-value < 0.05) specific for each major cluster after normalization, relative to Fig. 6E. (C) The fraction of subpopulations in total cells (cluster 0-7). (D) UMAP feature plots showing expression distribution of cell type specific genes in cell subpopulations, including RGL marker genes (VIM, HOPX, LPAR1 and SOX2), neurogenic development genes (STMN1), and reactive astrocytes marker gene (OSMR, TIMP and LGALS3). (E) The dynamic expression of cell type specific genes along the pseudotime. Each dot represents an individual cell. These representative genes included RGL genes PAX6 and HOPX, reactivated NSC genes VIM, CD44, TNC, CHI3L1 and SOX2, and cell cycle genes CKAP5 and RANGAP1, and neuroblast gene STMN2. Related to Figure 6.

**Figure 6-figure supplement 11. Integration of our snRNA-seq dataset with other published data.** (A) ETNPPL as a new NSC marker and STMN1/STMN2 as new immature neuron markers validated in Wang's study were verified in our study. (B) Integration of Zhou's snRNA-seq dataset of 14 aged donors (from 60-92 years old) with our snRNA-seq dataset. We did not detect evident pNSC, aNSC or NB populations in their dataset (circle with a dotted line). (C and D) UMAP visualization of pNSC/aNSC markers (TNC and VIM) (C), and neuroblast markers (STMN1 and NRGN) (D) in our and Zhou's snRNA-seq dataset. Related to discussion. (E to G) The snRNA-seq data set from Wang et al (Cell Research, 2022a) (E), Franjic et al. (Neuron 2022) (F), and Ayhan et al. (Neuron 2021) (G) were mapped onto our snRNA-seq data set using the "multimodal reference mapping" method. Based on the mapping analysis, astrocytes, qNSCs, aNSCs and NB were identified with varying correlation efficiencies in different datasets. The color scale represents the predicted correlation score.

### Supplement tables

**Figure 1-source data 1**. Patient information and the expression of findmarker genes used to identify cell populations in UMAP. Related to **Figure 1**
**Figure 2-source data 2.** The differential expression genes and relate GO terms of aNSC compared with pNSC.

**Figure 3-source data 3**. Potential marker genes identified by Findallmarker and scHPF. Related to **Figure 3**.
**Figure 4-source data 4.** Genes and enriched GO terms of Figure 4B and 4F. Related to **Figure 4**.
**Figure 5-source data 5.** Genes and enriched GO terms of qNSC1, qNSC2, pNSC, aNSC and NB populations during aging. Related to **Figure 5**.
**Figure 6-source data 6.** Genes and enriched GO terms of Figure 6F, 6H and Figure 6-S9A. Related to **Figure 6**.

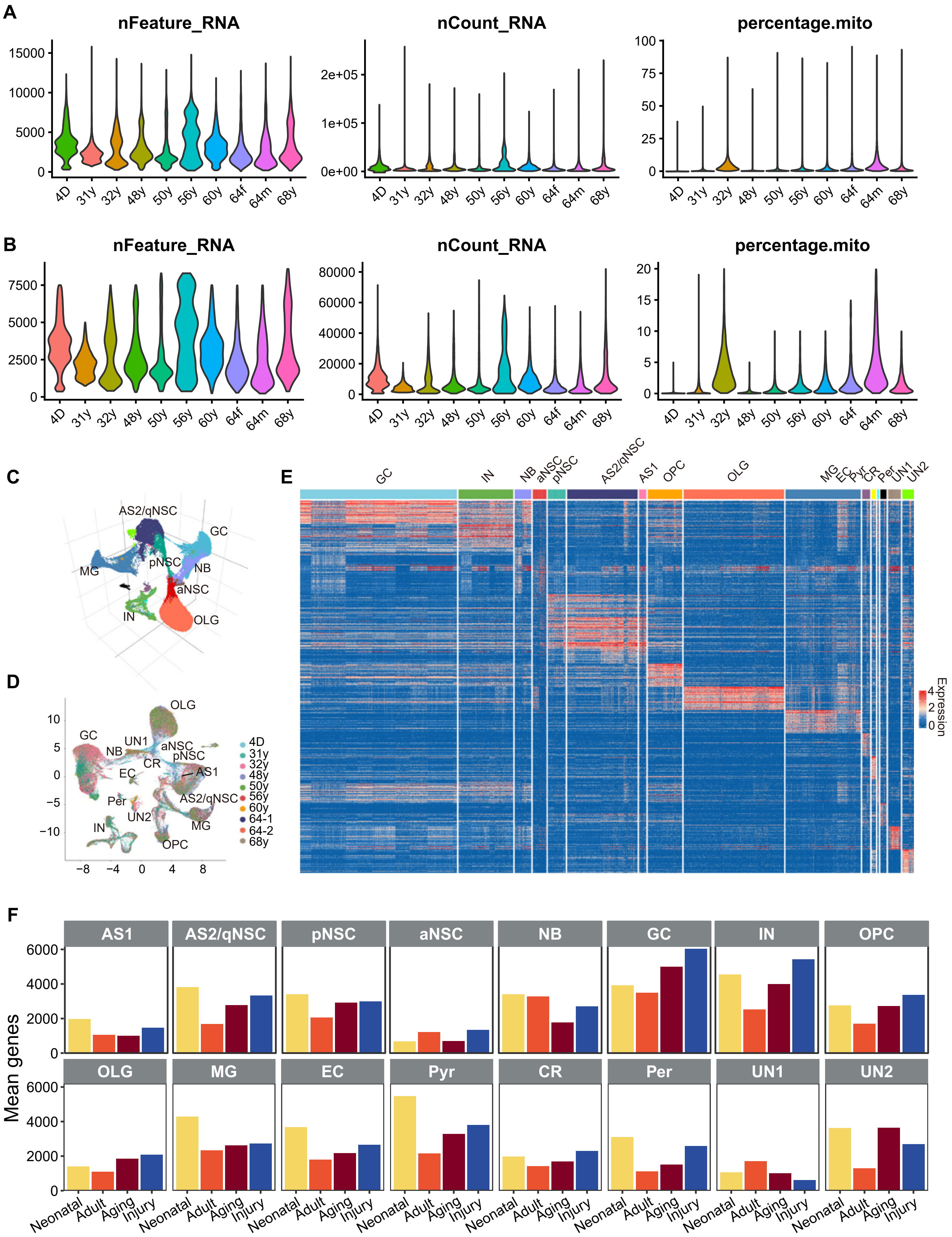

A
nFeature_RNA
nCount_RNA
percentage.mito
4D 31y 32y 48y 50y 56y 60y 64f 64m 68y
B
nFeature_RNA
nCount_RNA
percentage.mito
C
AS2/qNSC
GC
MG
pNSC
NB
aNSC
IN
OLG
D
OLG
GC
UN1
NB
aNSC
pNSC
CR
EC
AS1
AS2/qNSC
Per
UN2
IN
MG
OPC
4D
31y
32y
48y
50y
56y
60y
64-1
64-2
68y
E
GC IN NB aNSC pNSC AS2/qNSC AS1 OPC OLG MG EC Pyr CR Per UN1 UN2
Expression
F
AS1 AS2/qNSC pNSC aNSC NB GC IN OPC
OLG MG EC Pyr CR Per UN1 UN2
Mean genes
Neonatal Adult Aging Injury

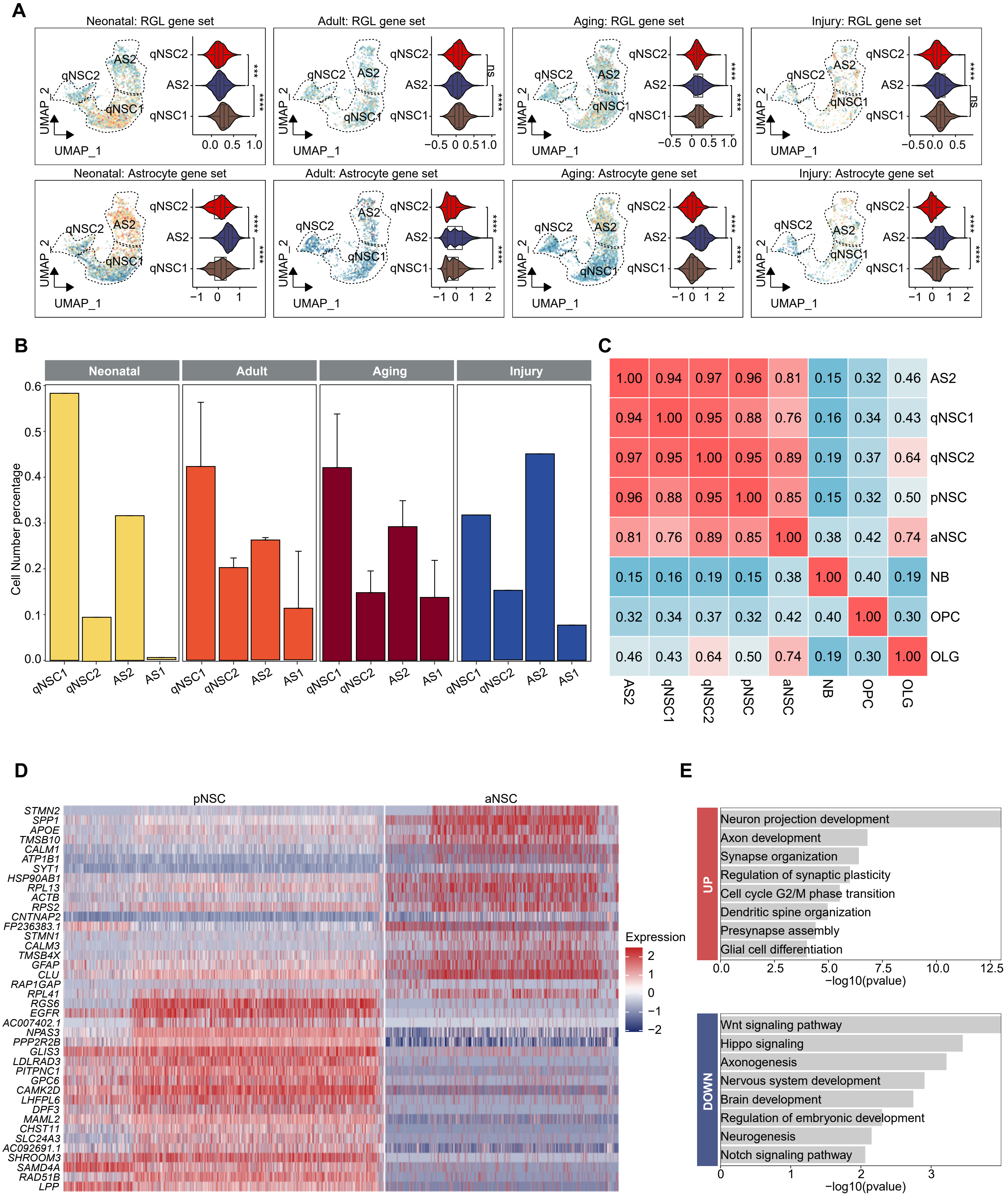
A
Neonatal: RGL gene set
Adult: RGL gene set
Aging: RGL gene set
Injury: RGL gene set
Neonatal: Astrocyte gene set
Adult: Astrocyte gene set
Aging: Astrocyte gene set
Injury: Astrocyte gene set
qNSC2
AS2
qNSC1
UMAP_1
UMAP_2
B
Neonatal
Adult
Aging
Injury
Cell Number percentage
qNSC1
qNSC2
AS2
AS1
C
AS2
qNSC1
qNSC2
pNSC
aNSC
NB
OPC
OLG
D
pNSC
aNSC
Expression
E
UP
Neuron projection development
Axon development
Synapse organization
Regulation of synaptic plasticity
Cell cycle G2/M phase transition
Dendritic spine organization
Presynapse assembly
Glial cell differentiation
−log10(pvalue)
DOWN
Wnt signaling pathway
Hippo signaling
Axonogenesis
Nervous system development
Brain development
Regulation of embryonic development
Neurogenesis
Notch signaling pathway
−log10(pvalue)

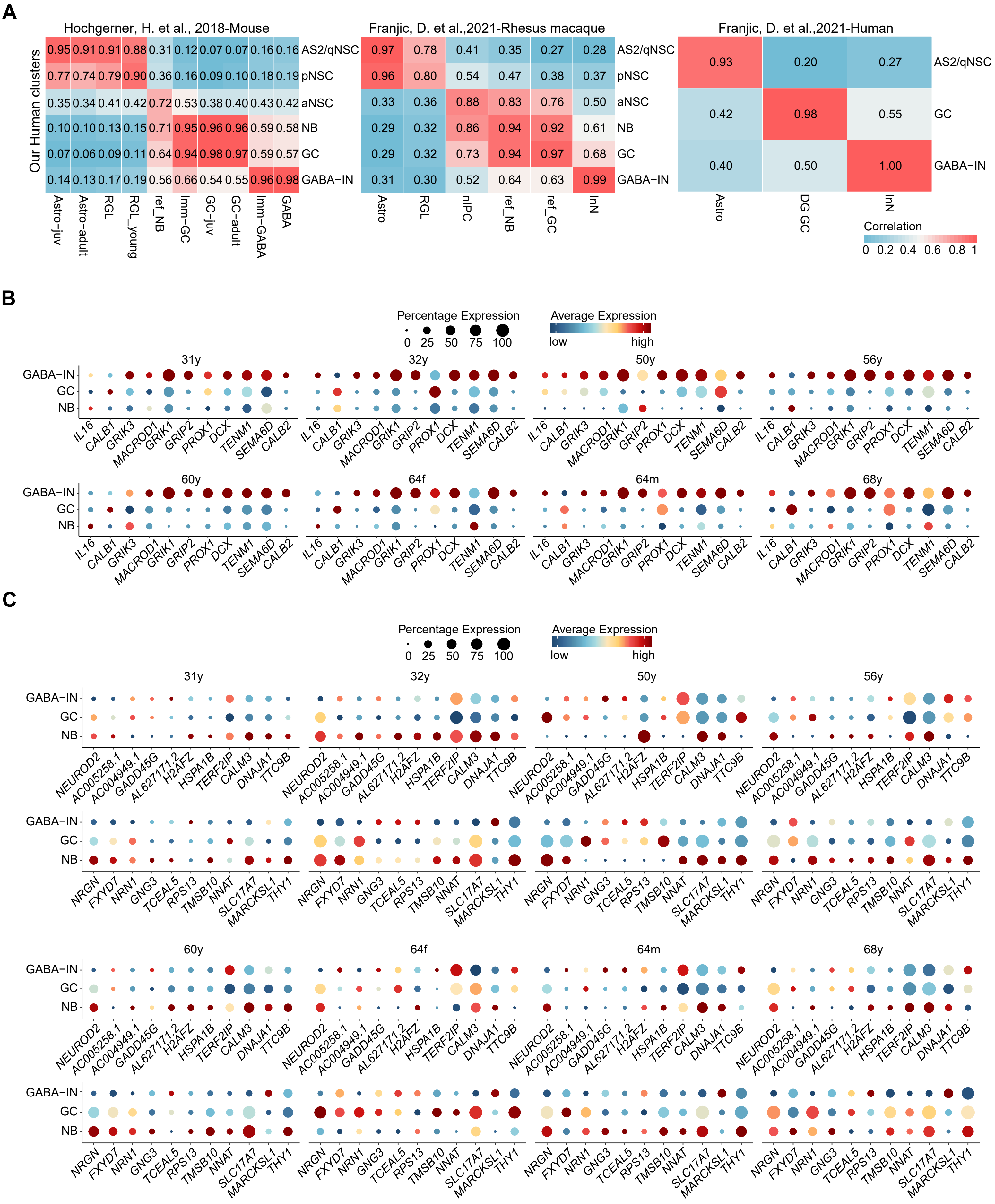
A
Hochgerner, H. et al., 2018-Mouse
Franjic, D. et al.,2021-Rhesus macaque
Franjic, D. et al.,2021-Human
Our Human clusters
0.95 0.91 0.91 0.88 0.31 0.12 0.07 0.07 0.16 0.16 AS2/qNSC
0.77 0.74 0.79 0.90 0.36 0.16 0.09 0.10 0.18 0.19 pNSC
0.35 0.34 0.41 0.42 0.72 0.53 0.38 0.40 0.43 0.42 aNSC
0.10 0.10 0.13 0.15 0.71 0.95 0.96 0.96 0.59 0.58 NB
0.07 0.06 0.09 0.11 0.64 0.94 0.98 0.97 0.59 0.57 GC
0.14 0.13 0.17 0.19 0.56 0.66 0.54 0.55 0.96 0.98 GABA−IN
Astro−juv Astro−adult RGL RGL_young ref_NB Imm−GC GC−juv GC−adult Imm−GABA GABA
0.97 0.78 0.41 0.35 0.27 0.28 AS2/qNSC
0.96 0.80 0.54 0.47 0.38 0.37 pNSC
0.33 0.36 0.88 0.83 0.76 0.50 aNSC
0.29 0.32 0.86 0.94 0.92 0.61 NB
0.29 0.32 0.73 0.94 0.97 0.68 GC
0.31 0.30 0.52 0.64 0.63 0.99 GABA−IN
Astro RGL nIPC ref_NB ref_GC InN
0.93 0.20 0.27 AS2/qNSC
0.42 0.98 0.55 GC
0.40 0.50 1.00 GABA−IN
Astro DG GC InN
Correlation
0 0.2 0.4 0.6 0.8 1
B
Percentage Expression
0 25 50 75 100
Average Expression
low high
31y 32y 50y 56y 60y 64f 64m 68y
GABA−IN
GC
NB
IL16 CALB1 GRIK3 MACROD1 GRIK1 GRIP2 PROX1 DCX TENM1 SEMA6D CALB2
C
Percentage Expression
0 25 50 75 100
Average Expression
low high
31y 32y 50y 56y 60y 64f 64m 68y
GABA−IN
GC
NB
NEUROD2 AC005258.1 AC004949.1 GADD45G AL627171.2 H2AFZ HSPA1B TERF2IP CALM3 DNAJA1 TTC9B
NRGN FXYD7 NRN1 GNG3 TCEAL5 RPS13 TMSB10 NNAT SLC17A7 MARCKSL1 THY1

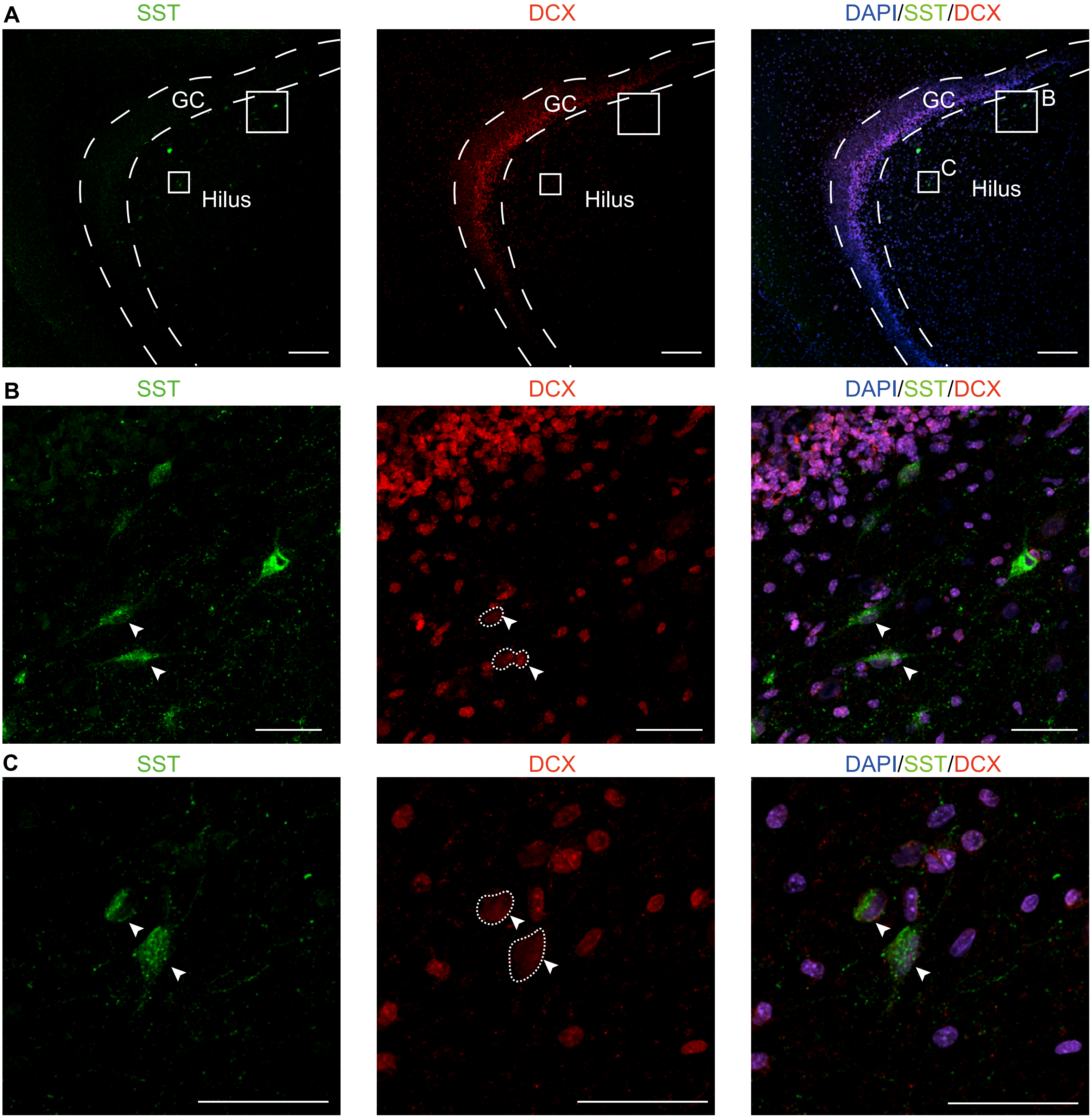
A
SST
DCX
DAPI/SST/DCX
GC
Hilus
B
C
SST
DCX
DAPI/SST/DCX
C
SST
DCX
DAPI/SST/DCX

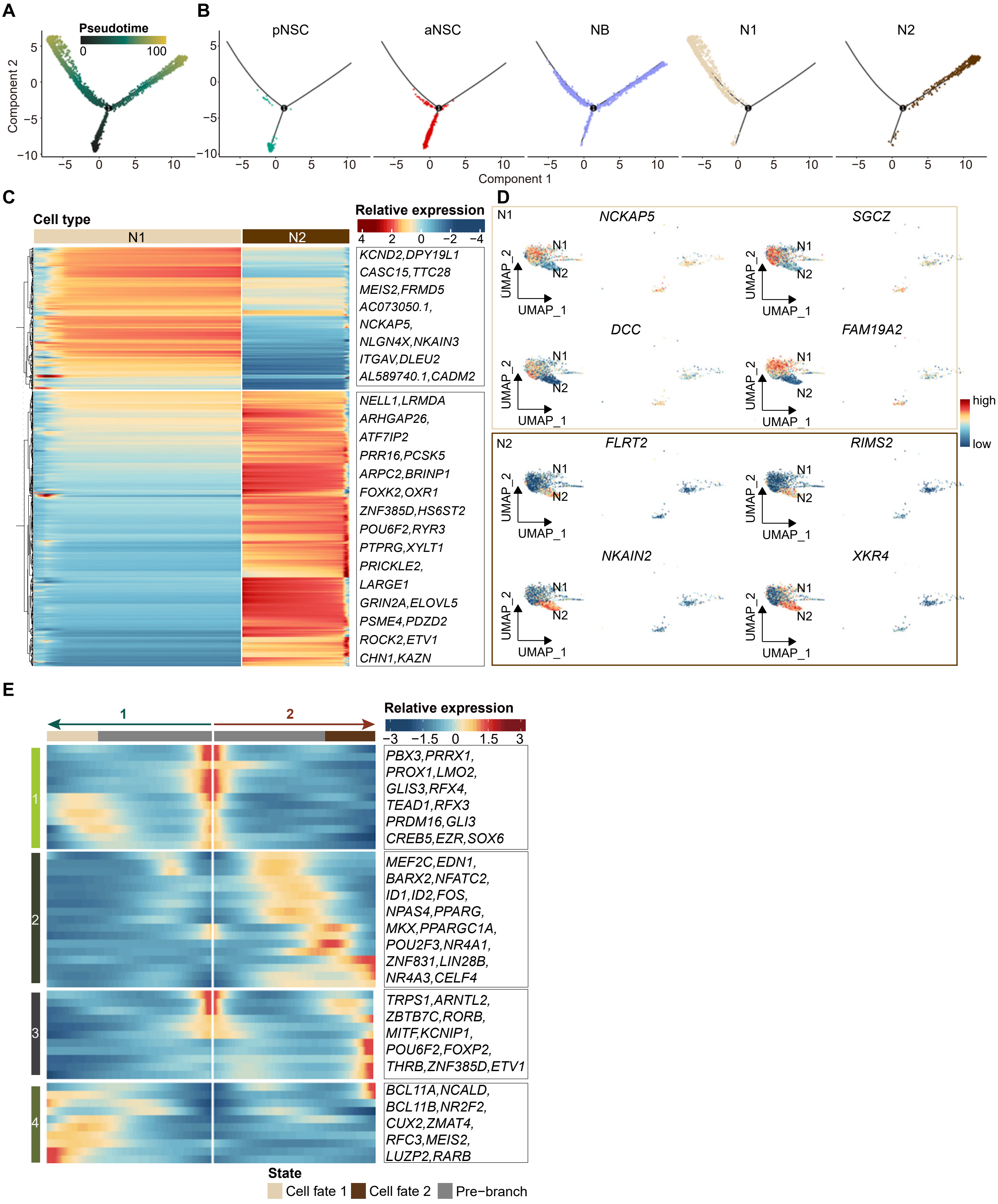

A
Pseudotime
0
100
Component 2
B
pNSC
aNSC
NB
N1
N2
Component 1
C
Cell type
N1
N2
Relative expression
KCND2,DPY19L1
CASC15,TTC28
MEIS2,FRMD5
AC073050.1,
NCKAP5,
NLGN4X,NKAIN3
ITGAV,DLEU2
AL589740.1,CADM2
NELL1,LRMDA
ARHGAP26,
ATF7IP2
PRR16,PCSK5
ARPC2,BRINP1
FOXK2,OXR1
ZNF385D,HS6ST2
POU6F2,RYR3
PTPRG,XYLT1
PRICKLE2,
LARGE1
GRIN2A,ELOVL5
PSME4,PDZD2
ROCK2,ETV1
CHN1,KAZN
D
N1
NCKAP5
SGCZ
DCC
FAM19A2
N2
FLRT2
RIMS2
NKAIN2
XKR4
UMAP_1
UMAP_2
high
low
E
Relative expression
PBX3,PRRX1,
PROX1,LMO2,
GLIS3,RFX4,
TEAD1,RFX3
PRDM16,GLI3
CREB5,EZR,SOX6
MEF2C,EDN1,
BARX2,NFATC2,
ID1,ID2,FOS,
NPAS4,PPARG,
MKX,PPARGC1A,
POU2F3,NR4A1,
ZNF831,LIN28B,
NR4A3,CELF4
TRPS1,ARNTL2,
ZBTB7C,RORB,
MITF,KCNIP1,
POU6F2,FOXP2,
THRB,ZNF385D,ETV1
BCL11A,NCALD,
BCL11B,NR2F2,
CUX2,ZMAT4,
RFC3,MEIS2,
LUZP2,RARB
State
Cell fate 1
Cell fate 2
Pre−branch

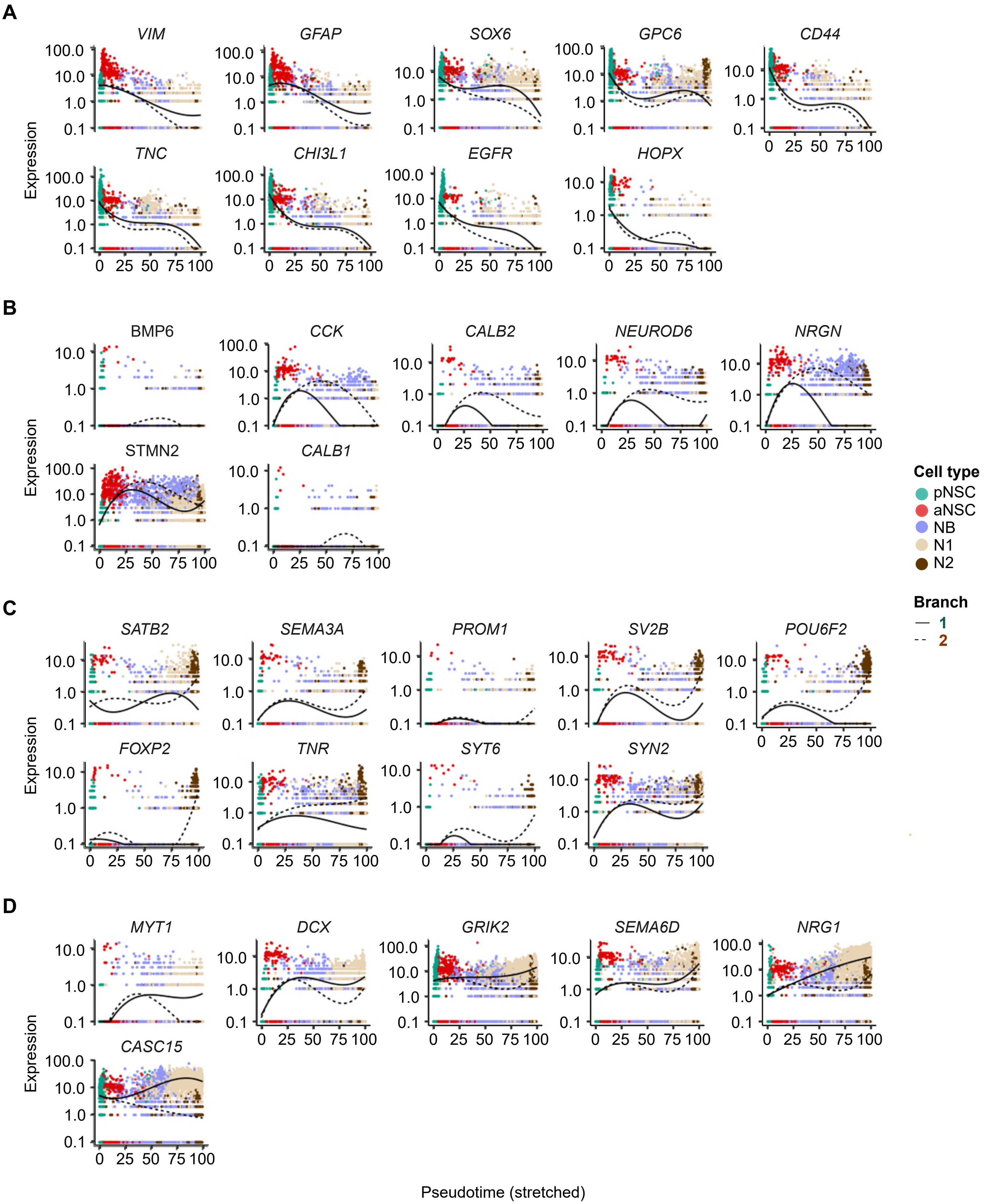

A
VIM
GFAP
SOX6
GPC6
CD44
TNC
CHI3L1
EGFR
HOPX
B
BMP6
CCK
CALB2
NEUROD6
NRGN
STMN2
CALB1
C
SATB2
SEMA3A
PROM1
SV2B
POU6F2
FOXP2
TNR
SYT6
SYN2
D
MYT1
DCX
GRIK2
SEMA6D
NRG1
CASC15
Expression
Pseudotime (stretched)
Cell type
pNSC
aNSC
NB
N1
N2
Branch
1
2

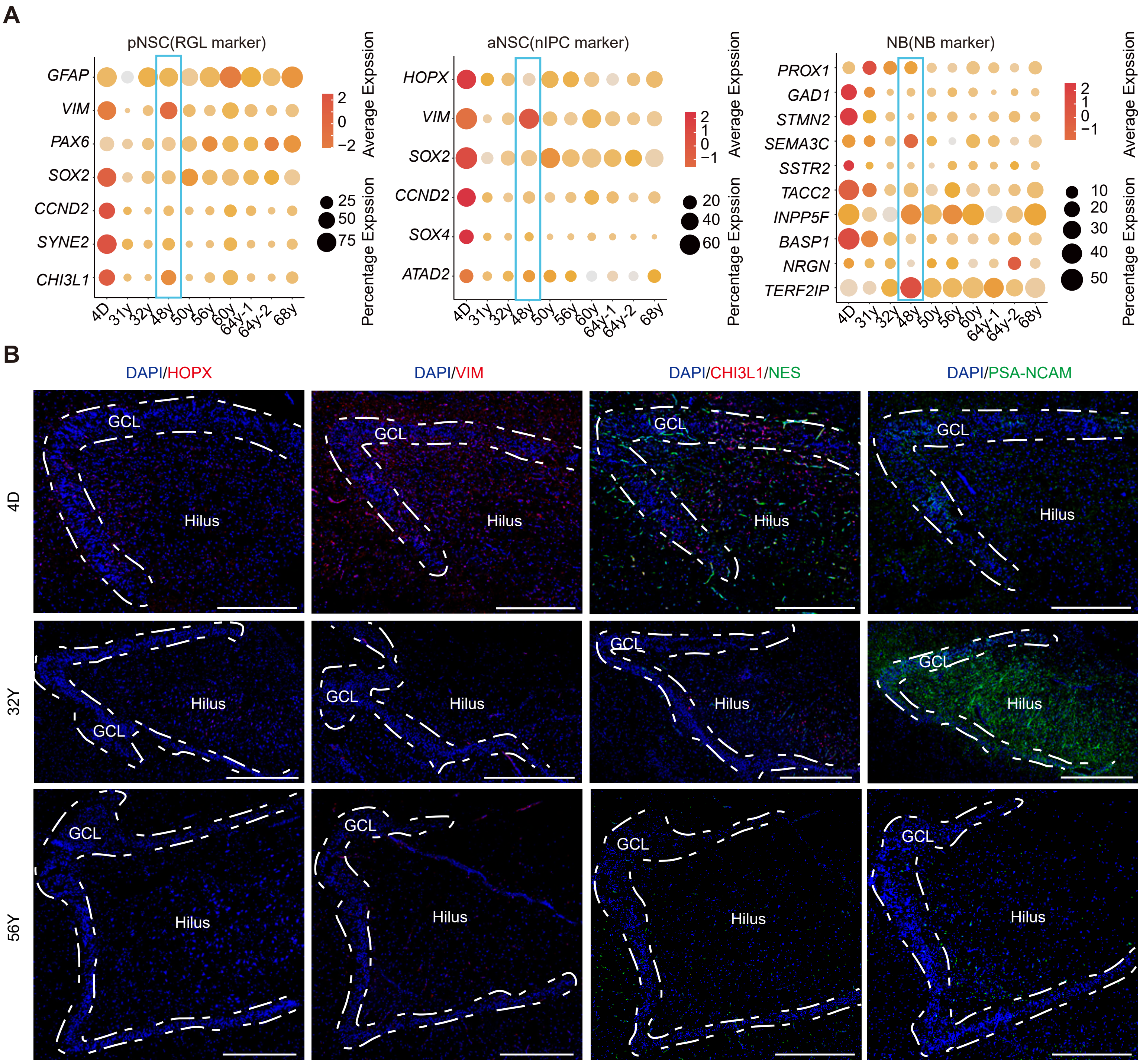

A
pNSC(RGL marker)
aNSC(nIPC marker)
NB(NB marker)
GFAP
VIM
PAX6
SOX2
CCND2
SYNE2
CHI3L1
HOPX
SOX4
ATAD2
PROX1
GAD1
STMN2
SEMA3C
SSTR2
TACC2
INPP5F
BASP1
NRGN
TERF2IP
4D
31y
32y
48y
50y
56y
60y
64y-1
64y-2
68y
Average Expssion
Percentage Expssion
B
DAPI/HOPX
DAPI/VIM
DAPI/CHI3L1/NES
DAPI/PSA-NCAM
4D
32Y
56Y
GCL
Hilus

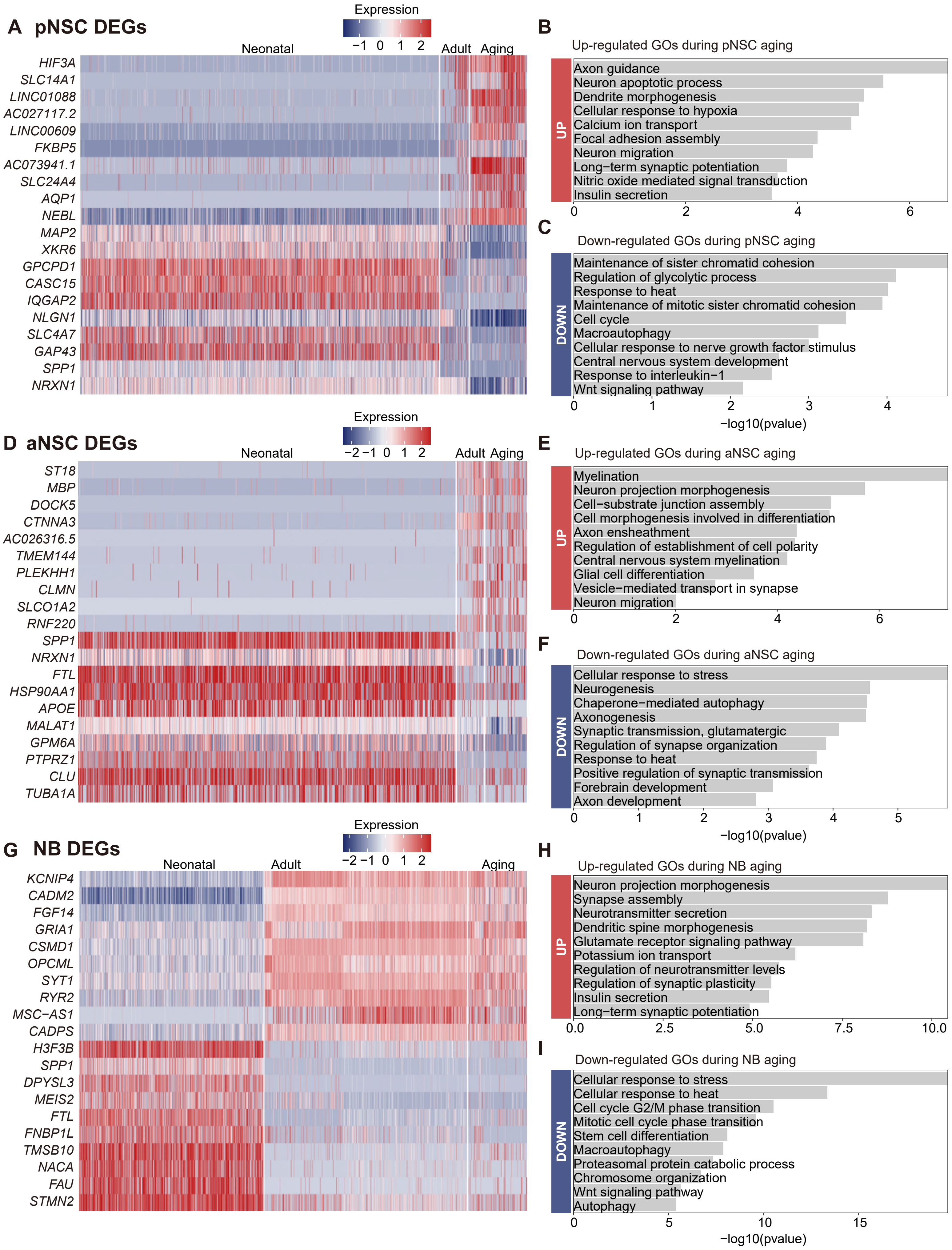
A pNSC DEGs
Expression
−1 0 1 2
Neonatal
Adult
Aging
HIF3A
SLC14A1
LINC01088
AC027117.2
LINC00609
FKBP5
AC073941.1
SLC24A4
AQP1
NEBL
MAP2
XKR6
GPCPD1
CASC15
IQGAP2
NLGN1
SLC4A7
GAP43
SPP1
NRXN1
B
Up-regulated GOs during pNSC aging
UP
Axon guidance
Neuron apoptotic process
Dendrite morphogenesis
Cellular response to hypoxia
Calcium ion transport
Focal adhesion assembly
Neuron migration
Long−term synaptic potentiation
Nitric oxide mediated signal transduction
Insulin secretion
0 2 4 6
C
Down-regulated GOs during pNSC aging
DOWN
Maintenance of sister chromatid cohesion
Regulation of glycolytic process
Response to heat
Maintenance of mitotic sister chromatid cohesion
Cell cycle
Macroautophagy
Cellular response to nerve growth factor stimulus
Central nervous system development
Response to interleukin−1
Wnt signaling pathway
0 1 2 3 4
−log10(pvalue)
D aNSC DEGs
Expression
−2 −1 0 1 2
Neonatal
Adult
Aging
ST18
MBP
DOCK5
CTNNA3
AC026316.5
TMEM144
PLEKHH1
CLMN
SLCO1A2
RNF220
SPP1
NRXN1
FTL
HSP90AA1
APOE
MALAT1
GPM6A
PTPRZ1
CLU
TUBA1A
E
Up-regulated GOs during aNSC aging
UP
Myelination
Neuron projection morphogenesis
Cell−substrate junction assembly
Cell morphogenesis involved in differentiation
Axon ensheathment
Regulation of establishment of cell polarity
Central nervous system myelination
Glial cell differentiation
Vesicle−mediated transport in synapse
Neuron migration
0 2 4 6
F
Down-regulated GOs during aNSC aging
DOWN
Cellular response to stress
Neurogenesis
Chaperone−mediated autophagy
Axonogenesis
Synaptic transmission, glutamatergic
Regulation of synapse organization
Response to heat
Positive regulation of synaptic transmission
Forebrain development
Axon development
0 1 2 3 4 5
−log10(pvalue)
G NB DEGs
Expression
−2 −1 0 1 2
Neonatal
Adult
Aging
KCNIP4
CADM2
FGF14
GRIA1
CSMD1
OPCML
SYT1
RYR2
MSC−AS1
CADPS
H3F3B
SPP1
DPYSL3
MEIS2
FTL
FNBP1L
TMSB10
NACA
FAU
STMN2
H
Up-regulated GOs during NB aging
UP
Neuron projection morphogenesis
Synapse assembly
Neurotransmitter secretion
Dendritic spine morphogenesis
Glutamate receptor signaling pathway
Potassium ion transport
Regulation of neurotransmitter levels
Regulation of synaptic plasticity
Insulin secretion
Long−term synaptic potentiation
0.0 2.5 5.0 7.5 10.0
I
Down-regulated GOs during NB aging
DOWN
Cellular response to stress
Cellular response to heat
Cell cycle G2/M phase transition
Mitotic cell cycle phase transition
Stem cell differentiation
Macroautophagy
Proteasomal protein catabolic process
Chromosome organization
Wnt signaling pathway
Autophagy
0 5 10 15
−log10(pvalue)

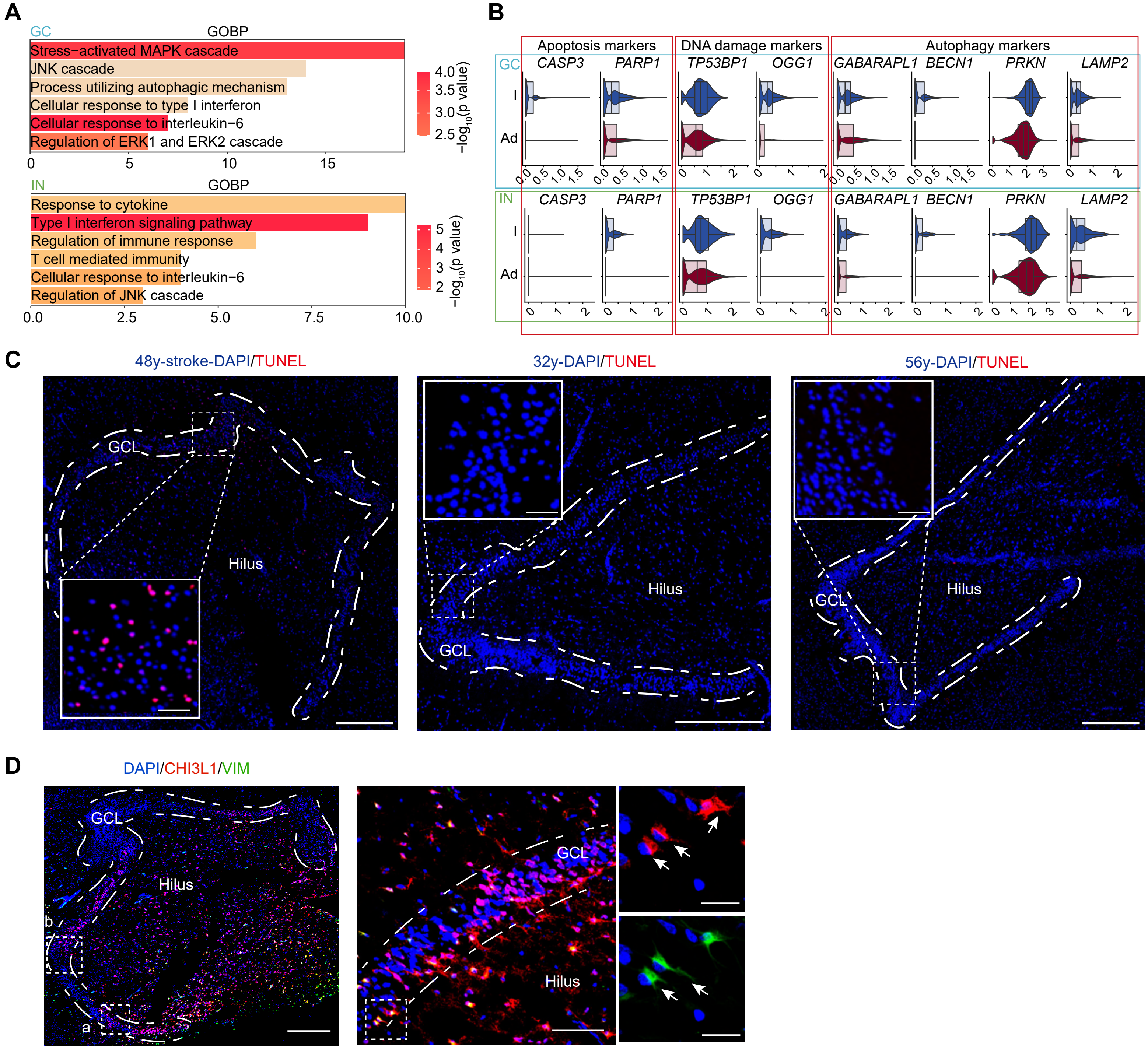

A
GC
GOBP
Stress−activated MAPK cascade
JNK cascade
Process utilizing autophagic mechanism
Cellular response to type I interferon
Cellular response to interleukin−6
Regulation of ERK1 and ERK2 cascade
−log10(p value)
IN
GOBP
Response to cytokine
Type I interferon signaling pathway
Regulation of immune response
T cell mediated immunity
Cellular response to interleukin−6
Regulation of JNK cascade
B
Apoptosis markers
DNA damage markers
Autophagy markers
CASP3
PARP1
TP53BP1
OGG1
GABARAPL1
BECN1
PRKN
LAMP2
I
Ad
C
48y-stroke-DAPI/TUNEL
32y-DAPI/TUNEL
56y-DAPI/TUNEL
GCL
Hilus
D
DAPI/CHI3L1/VIM
GCL
Hilus
a
b

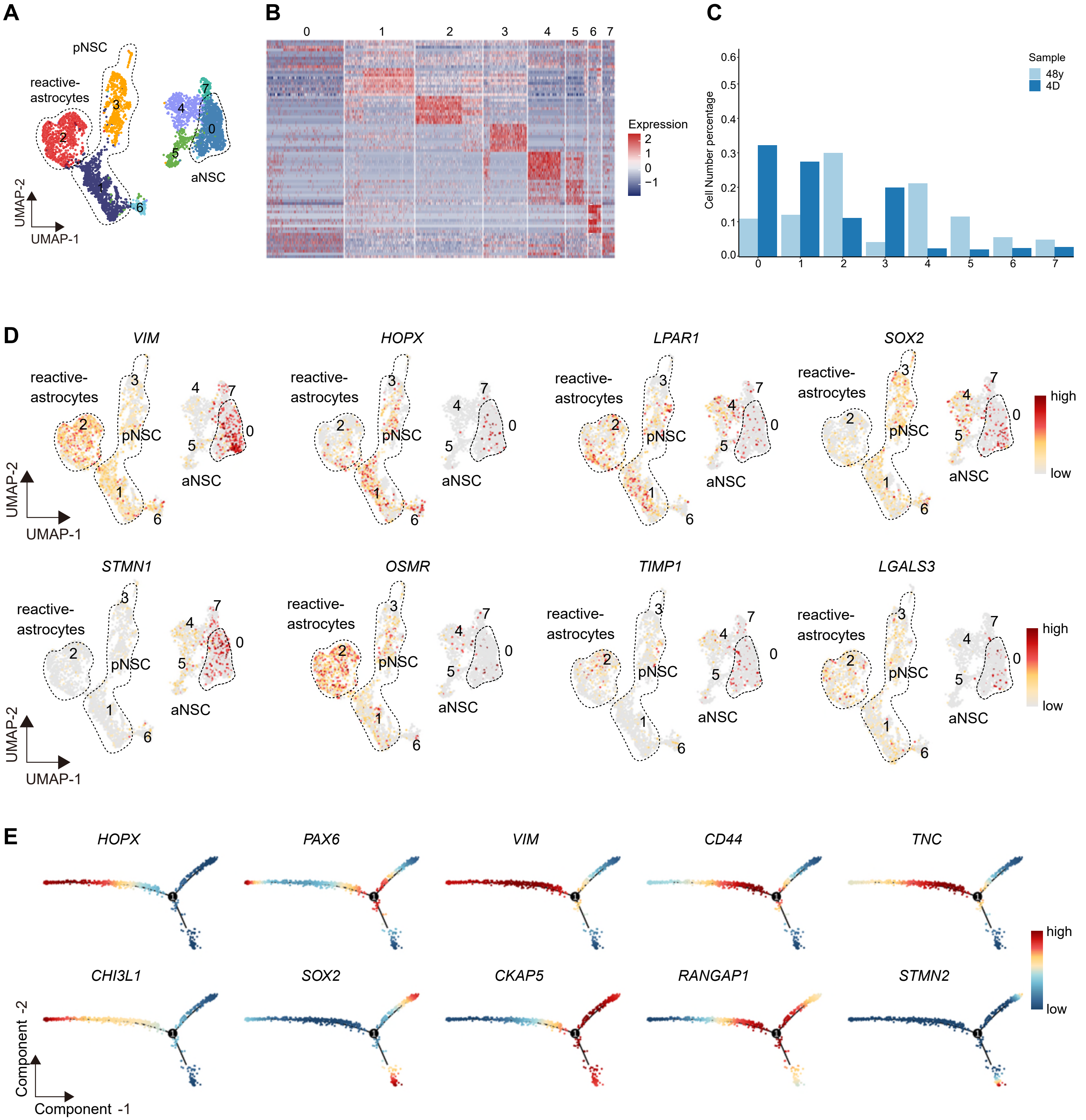

A
pNSC
reactive-
astrocytes
aNSC
UMAP-2
UMAP-1
B
Expression
C
Sample
48y
4D
Cell Number percentage
D
VIM
HOPX
LPAR1
SOX2
STMN1
OSMR
TIMP1
LGALS3
high
low
E
PAX6
CD44
TNC
CHI3L1
CKAP5
RANGAP1
STMN2
Component -2
Component -1

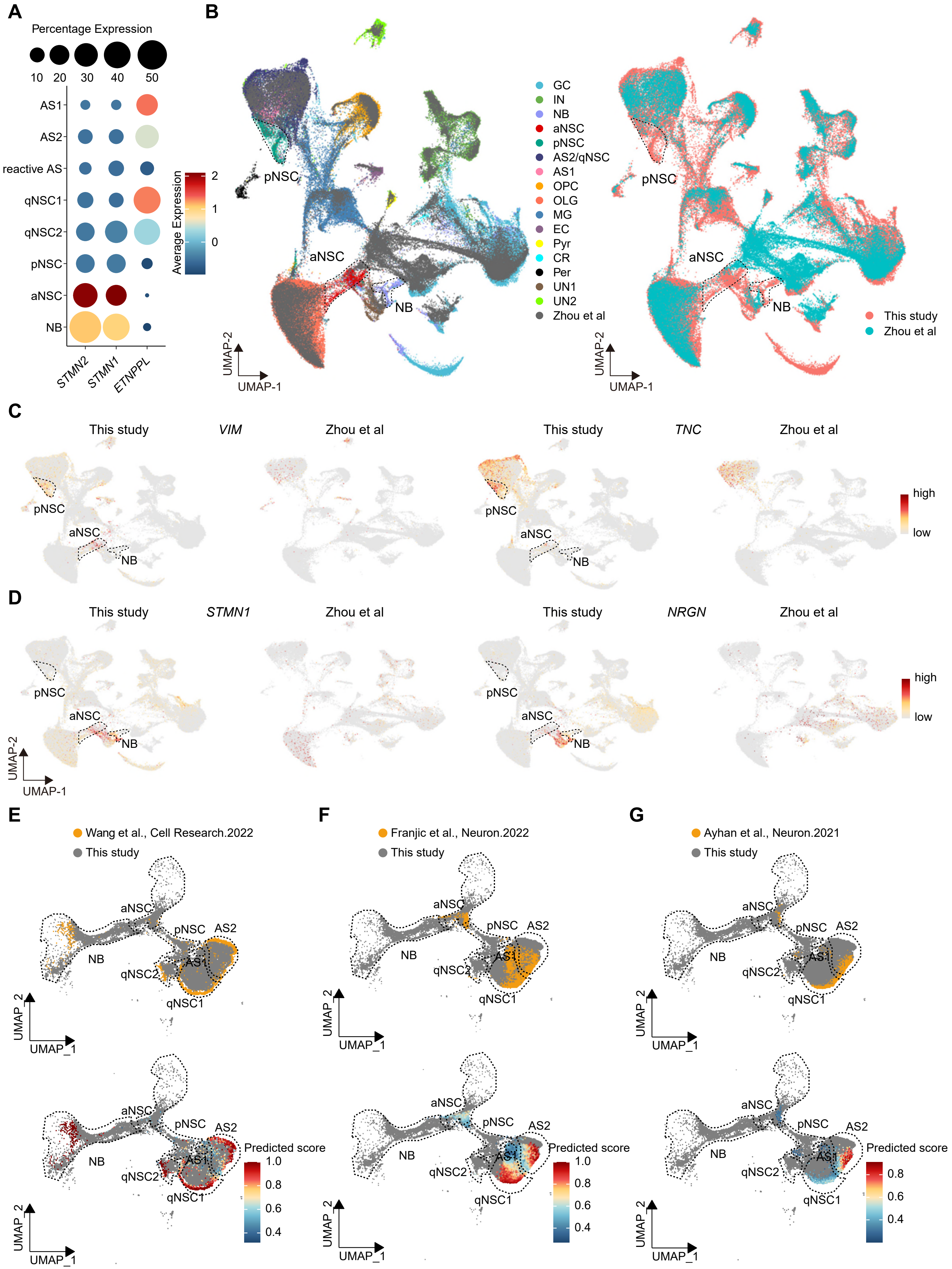

A
Percentage Expression
10 20 30 40 50
AS1
AS2
reactive AS
qNSC1
qNSC2
pNSC
aNSC
NB
STMN2
STMN1
ETNPPL
Average Expression
2
1
0
B
GC
IN
NB
aNSC
pNSC
AS2/qNSC
AS1
OPC
OLG
MG
EC
Pyr
CR
Per
UN1
UN2
Zhou et al
This study
Zhou et al
UMAP-2
UMAP-1
C
This study
VIM
Zhou et al
TNC
high
low
D
STMN1
NRGN
E
Wang et al., Cell Research.2022
This study
F
Franjic et al., Neuron.2022
G
Ayhan et al., Neuron.2021
Predicted score
UMAP_2
UMAP_1
